# Preclinical Multi-Omic Assessment of Pioglitazone in Skeletal Muscles of Mice Implanted with Human HER2/neu Overexpressing Breast Cancer Xenografts

**DOI:** 10.1101/2024.04.15.589557

**Authors:** Stuart A. Clayton, Alan D. Mizener, Marcella Whetsell, Lauren E. Rentz, Ethan Meadows, Werner Geldenhuys, Emidio E. Pistilli

## Abstract

Breast cancer (BC) is the most prevalent cancer worldwide and is accompanied by fatigue during both active disease and remission in the majority of cases. Our lab has measured fatigue in isolated muscles from treatment-naive BC patient-derived orthotopic xenograft (BC-PDOX) mice. Here, we conducted a preclinical trial of pioglitazone in BC-PDOX mice to determine its efficacy in ameliorating BC-induced muscle fatigue, as well as its effects on transcriptomic, metabolomic, and lipidomic profiles in skeletal muscle.

**Methods:** The pioglitazone and vehicle groups were treated orally for 4 weeks upon reaching a tumor volume of 600 mm^3^. Whole-animal indirect calorimetry was used to evaluate systemic metabolic states. The transcriptome was profiled using short-read bulk RNA sequencing (RNA-seq). Liquid chromatography-tandem mass spectrometry (LC-MS/MS) was used to profile the metabolome and lipidome. Fast and slow skeletal muscle function were evaluated using isolated *ex vivo* testing.

**Results:** Pioglitazone was associated with a significant overall decrease in metabolic rate, with no changes in substrate utilization. RNA-seq supported the downstream effects of pioglitazone on target genes and displayed considerable upregulation of mitochondrial bioenergetic pathways. Skeletal muscle metabolomic and lipidomic profiles exhibited dysregulation in response to BC, which was partially restored in pioglitazone-treated mice compared to vehicle-treated BC-PDOX mice. Despite molecular support for pioglitazone’s efficacy, isolated muscle function was not affected by pioglitazone treatment.

**Conclusions:** BC induces multi-omic dysregulation in skeletal muscle, which pioglitazone partially ameliorates. Future research should focus on profiling systemic metabolic dysfunction, identifying molecular biomarkers of fatigue, and testing alternative pioglitazone treatment regimens.

**Statement of Translational Relevance:** Breast cancer-induced fatigue is a prevalent and debilitating symptom that affects a majority of patients, leading to early treatment discontinuation and poorer outcomes. Despite its significant impact on patient quality of life, there are currently no approved therapies for this condition. Our previous work in the clinically relevant breast cancer patient-derived orthotopic xenograft (BC-PDOX) mouse model suggests that disruptions in the PPARγ signaling pathway may contribute to the development of cancer-related fatigue. Using this model that recapitulates the fatigue phenotype observed in patients, we conducted a preclinical trial evaluating the FDA-approved PPARγ agonist, pioglitazone, as a treatment for fatigue. Our multi-omic analysis of skeletal muscle from BC-PDOX mice revealed that pioglitazone treatment partially restored dysregulated lipid profiles and mitochondrial bioenergetic transcriptomic alterations. These findings suggest that pioglitazone may have potential as a therapeutic option for managing cancer-related fatigue in breast cancer patients.

## Introduction

As of 2020, breast cancer (BC) is the most prevalent cancer globally, with over two million women diagnosed in that year (1). BC incidence continues to increase worldwide, with the highest rates occurring in transitioned countries, whereas the highest mortalities occur in transitioning countries (1). In the United States, the 43% decrease in mortality since 1989 (2) can be attributed to enhanced screening, better diagnostics, and advanced treatments (3). BC is a systemic disease and one of the most prevalent symptoms in BC patients is fatigue, with BC-associated skeletal muscle (SkM) fatigue being reported to affect between 62% and 85% of patients undergoing active treatment (4) and up to 66% of disease-free survivors (5). In contrast to these high rates of fatigue, the estimated cachexia rate in BC patients is one of the lowest of all cancer types (6), although rates can vary depending on the specific definition used. Fatigue persistence in disease-free cases emphasizes the need for effective treatments across disease stages, and the importance of studying the mechanistic underpinnings of BC-associated SkM fatigue. It has been suggested that BC patients with fatigue have worse outcomes and higher mortality (7). Despite the rates of fatigue, quality-of-life impact, and potential reduction in survival, there are currently no effective treatments to ameliorate this debilitating symptom.

The peroxisome proliferator-activated receptor (PPAR) family comprises three lipid-sensing nuclear transcription factors: PPARɑ, -β/δ, and -γ (8). Collectively, PPARs are pivotal regulators of mitochondrial fatty acid β-oxidation (9) (10), adipocyte differentiation and fat storage (11) (12), and insulin sensitivity (13). Our data suggest that disruption in PPAR signaling is a driver of fatigue using a BC patient-derived orthotopic xenograft (BC-PDOX) mouse model that recapitulates fatigue in the absence of cachexia (14) (15). Aberrations in PPAR signaling can lead to disruption of mitochondrial bioenergetics and lipid accumulation (16) (17) as well as transcriptome dysregulation (15).

Pioglitazone, a well-tolerated FDA-approved thiazolidinedione (TZD) and PPARγ-agonist, demonstrates a high affinity for both PPARγ isoforms (1 and 2) (18), which are expressed in both humans and mice (19) (20), and is indicated for the treatment of insulin resistance in type-2 diabetes mellitus (21). We have previously demonstrated that pioglitazone restores transcriptomic profiles in SkM of BC-PDOX mice and is implicated as a potential therapeutic option for treating BC-associated SkM fatigue (15). The aim of this study was to test the efficacy of pioglitazone treatment for four weeks as an intervention to reduce SkM fatigue in BC-PDOX mice implanted with HER2/neu overexpressing tumors. To explore the potential metabolic alterations that may contribute to fatigue, we also performed untargeted metabolomics and lipidomics. We hypothesized that treatment of BC-PDOX mice with pioglitazone for four weeks would demonstrate a reduction in fatigue of type II muscles, a rescued transcriptomic profile, and an overall decrease in metabolic rate compared to non-drug treated BC-PDOX mice.

## Materials and Methods

### Breast cancer patient-derived orthotopic xenograft (BC-PDOX) mouse model

We used 16 8-week old female NOD.Cg-*Prkdc*^*scid*^ *Il2rg*^*tm1Wjl*^/SzJ (NSG; RRID:IMSR_JAX:005557) mice obtained from Jackson Laboratory (The Jackson Laboratory, Bar Harbor, ME, USA). All NSG mice were genotyped using strain-specific probes to verify correct strain identity by Transnetyx (Transnetyx Inc., Cordova, TN, USA). Mice were housed at 22 °C in the AAALAC accredited vivarium at West Virginia University (WVU) on a standard 12:12 hr light:dark cycle in sterile polystyrene cages on soft-bedding and provided irradiated Tekklad (Inotiv, Maryland Heights, MO, USA) 18% protein rodent diet (3.1 kcal∙g^-1^) and Sulfatrim supplemented water (sulfamethoxazole, 0.26 mg∙mL^-1^; trimethoprim, 0.052 mg∙mL^-1^) Monday through Thursday and sterile water the remaining days, all *ad libitum*. HER2/neu overexpressing human tumor samples were obtained from the patient-derived xenograft (PDX) bank at WVU. Genetic comparability between the injected and original tumors was validated via short tandem repeat (STR) profiling (Supplementary Figure 1), and similarity was calculated using the following equation: % similarity = (# of Matching Alleles) / (# of Total Alleles Detected). To prepare PDXs for implantation, tumor fragments were minced and enzymatically dissociated using a Miltenyi Human Tumor Dissociation Kit and Miltenyi gentleMACS Octo Dissociator with Heaters (Miltenyi Biotec, Bergisch Gladbach, DE; RRID:SCR_020271). A single-cell suspension of a HER2/neu overexpressing PDX was injected bilaterally at the fourth inguinal nipple using a sterile ½ inch 26-gauge needle. Approximately 2 x 10^6^ cells were injected in 100 µL of a 1:1 (v/v) mixture of Cultrex basement membrane extract (BME) type 3 (Biotechne R&D Systems, Minneapolis, MN, USA) and sterile 1X phosphate buffered saline (PBS). Tumor volumes were monitored at least once per week by calipers until a composite volume of 200 mm^3^ was reached, followed by micro-ultrasound using Vevo F2 (VisualSonics, Bothell, WA, USA). Tumor 3D-volume was calculated from captured ultrasound images using Vevo LAB (version 5.8.2). Mice were euthanized upon completing treatment or earlier if they became moribund. Precise sample sizes used in each of the subsequent analyses are provided in Supplementary Table 1. All procedures followed the protocols approved by the WVU Institutional Animal Care and Use Committee (IACUC) and were conducted in accordance with the NIH guidelines for animal research.

**Figure 1.**
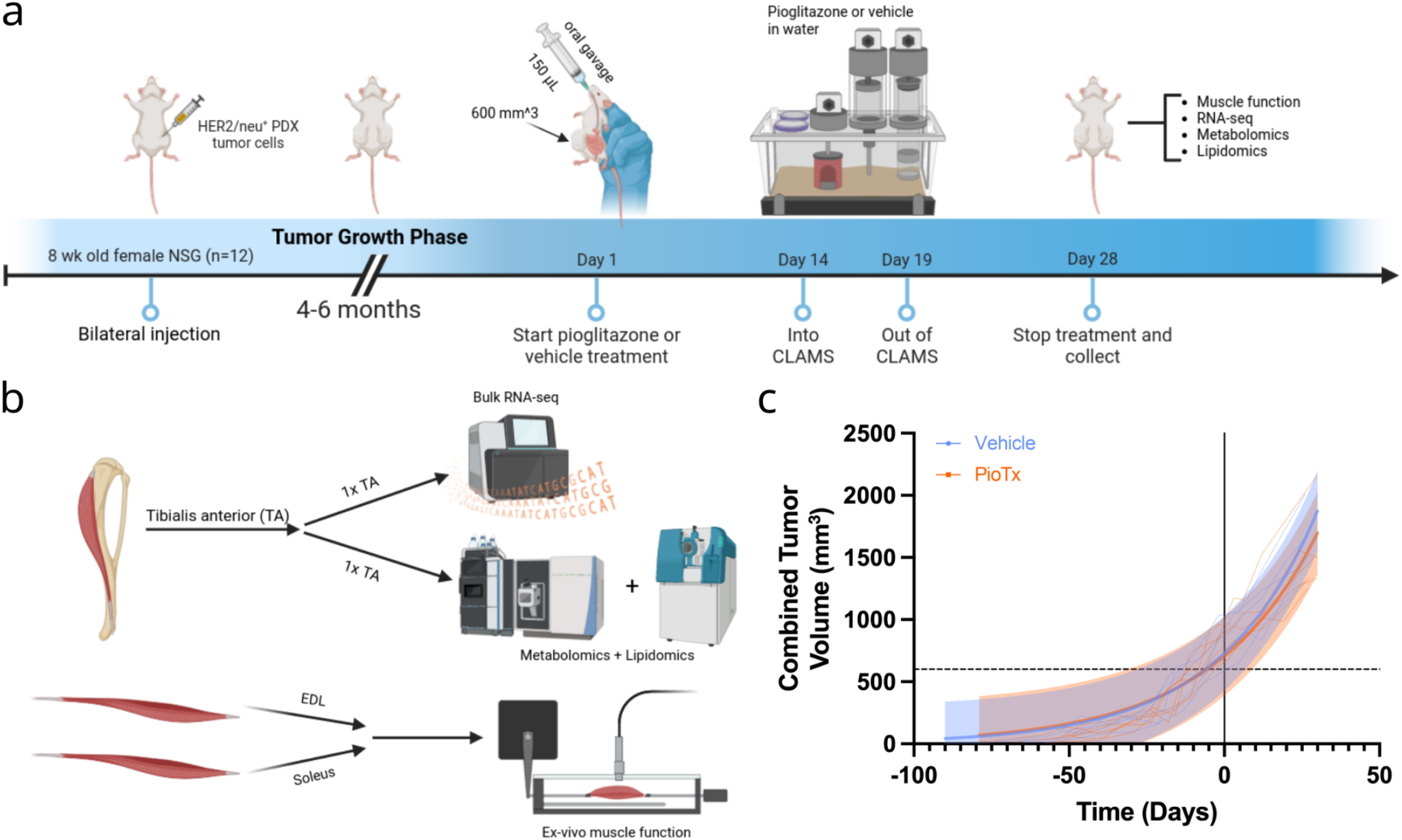
Overview and characterization of the BC-PDOX mouse model. (**a**) Experimental timeline. (**b**) Analysis of isolated muscles. One TA muscle was used for bulk RNA sequencing, while the contralateral was analyzed using metabolomics and lipidomics. The EDL and soleus muscles were used for *ex vivo* muscle function tests. (**c**) Exponential regression plot of longitudinal tumor volumes. Times for each volume measurement were adjusted to set the treatment start date to t = 0. Colored dashed lines represent combined tumor volume measurements for individual animals. Shaded regions depict the 95% prediction interval for each group. The black dashed line is at 600 mm^3^ for reference. (**a**,**b**) Created with Biorender.com. TA, tibialis anterior; EDL, extensor digitorum longus; CLAMS, comprehensive laboratory animal monitoring system; 1x TA refers to one TA muscle from each animal being used for bulk RNA-seq, and one being used for metabolomics/lipidomics.

### Pioglitazone preparation and dosing

BC-PDOX mice were randomized into either the pioglitazone- (PioTx) or vehicle-treated (vehicle) group. One mouse was euthanized prior to study completion because of tumor ulceration and was excluded from all analyses. Pioglitazone Hydrochloride (>98.0% purity) was purchased from TCI Chemicals (TCI Chemicals, Portland, OR, USA; Product #: P1901) and prepared at a dosage of 30 mg·kg^-1^·day^-1^, assuming an average body weight of 30 g, and dissolved by rocking for 1 hr at room temperature in 10% w/v sulfobutylether-β-cyclodextrin (Captisol, hereafter vehicle) (Captisol, San Diego, CA, USA). The drug was prepared weekly and was stored at 4 °C. Pioglitazone (0.9 mg per dose) or vehicle was thoroughly shaken to account for drug settling and was administered to all mice for at least 28 days. Drug administration began at a tumor volume of 600 mm^3^ and during study weeks 1, 2, and 4 was administered via daily oral gavage at 12 pm (± 52 min) in a 150 µL bolus using 38 mm silicone-tipped 18-gauge plastic gavage needles (Pet Surgical, Phoenix, AZ, USA). Drug administration during study week 3, while animals were monitored using indirect calorimetry, was administered via drinking water. Drug concentration was based on historical water consumption, and post-hoc drug dosage calculations yielded an average of 26 mg·kg^-1^·day^-1^ (18-30 mg·kg^-1^·day^-1^).

### Bulk RNA isolation and sequencing

Tibialis anterior (TA) muscles from PioTx and vehicle NSG mice were harvested, flash-frozen in liquid nitrogen, and stored at -80 °C until further processing. TA muscles were thawed and cut into pieces weighing ≤ 30 mg, followed by mechanical dissociation using a TissueRuptor (Qiagen, Venlo, Limburg, NL). Bulk RNA was isolated using the RNeasy Fibrous Tissue Mini Kit (Qiagen) according to the manufacturer’s protocol. A NanoDrop spectrophotometer was used to quantify RNA concentration (336.4 ± 209.3 ng∙µL^-1^) and purity using A260/280 values (2.11 ± 0.02). The isolated RNA was shipped to Admera Health (Admera Health, Plainfield, NJ, USA) on dry ice for library preparation and sequencing. RNA integrity was assessed using an RNA Tapestation (Agilent Technologies Inc., Santa Clara, CA, USA) and quantified by Qubit 2.0 RNA HS assay (ThermoFisher, Massachusetts, USA). RNA integrity numbers (RIN) were ≥ 7.7 (Supplementary Figure 2a).

**Figure 2.**
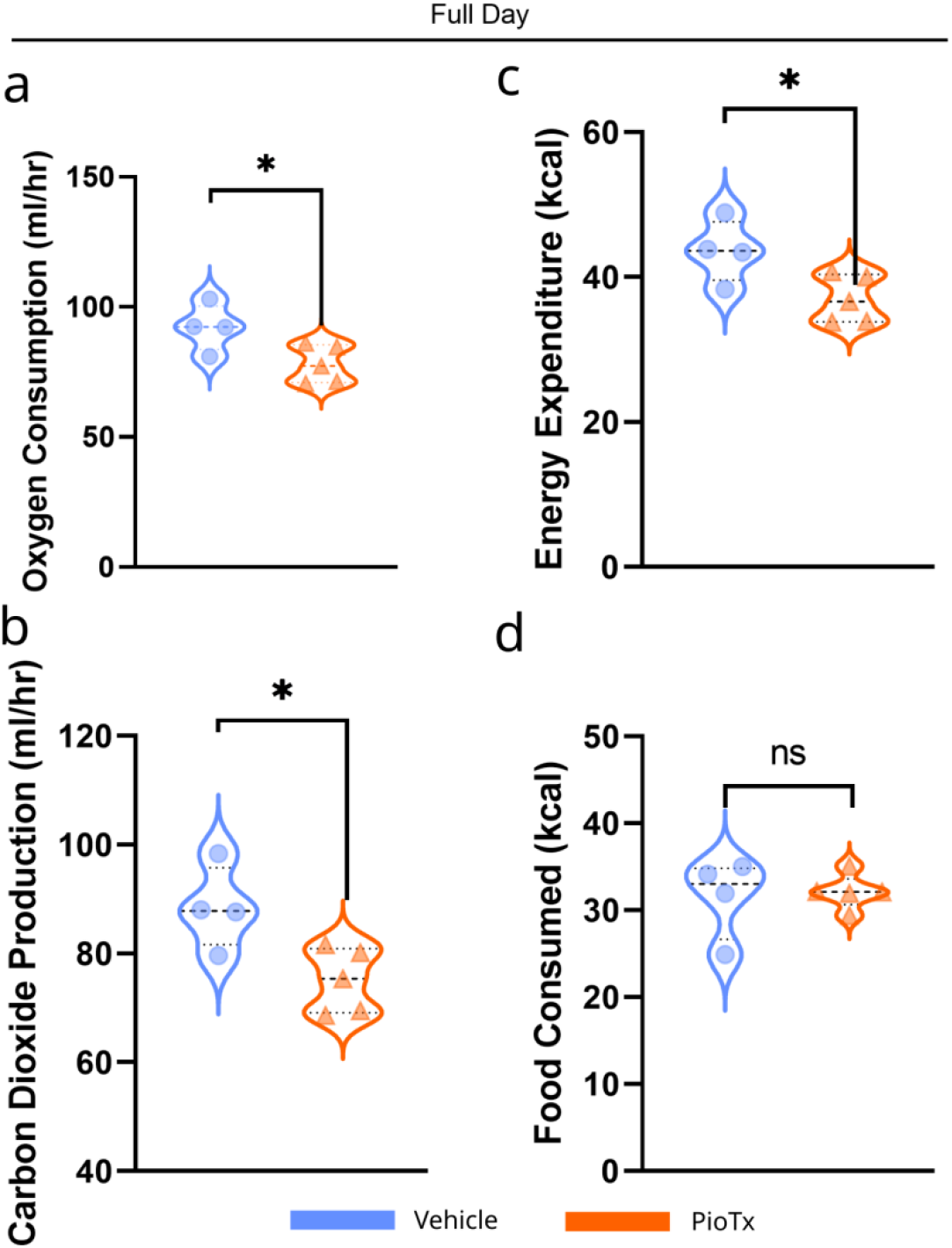
Whole-animal indirect calorimetry. Violin plots comparing metabolic variables among vehicle (blue, n=4) and PioTx (orange, n=5) mice. (**a**) Average oxygen consumption across the full day in ml∙hr^-1^. (**b**) Average carbon dioxide production across the full day in ml∙hr^-1^. (**c**) Cumulative energy expenditure (kcal) across the 96 hour period. (**d**) Total food consumed (kcal), across the 96 hour period. The energy density of the food provided was 3.1 kcal∙g^-1^. * < 0.05.

Poly(A) selection and cDNA libraries were constructed using the NEBNext Ultra II Directional RNA Library Prep Kit for Illumina (New England BioLabs Inc., Massachusetts, USA). The final library size was approximately 430 bp with an insert size of approximately 300 bp. Illumina 8-nt dual indices were used. Samples were pooled at equimolar concentrations and sequenced on an Illumina NovaSeq (Illumina, California, USA) with a read length configuration of 150 PE for 40M paired-end reads per sample (20M in each direction). (BioProject ID: PRJNA1076666 murine samples).

### Bulk-RNAseq data analysis

Paired reads had a 5’ trim of 13 bases and a 3’ trim of 87 bases, which were then aligned to the Ensembl GRCm39.109 mouse reference genome using HISAT2 (version 2.2.1; RRID:SCR_015530) (22). RNA samples (n=11) averaged 36,024,271 ± 10,109,476 reads per sample (Supplementary Figure 2b) with a 96.9 ± 0.21% mapping rate to the GRCm39.109 genome. Raw read counts were generated using the featureCounts function of Subread (version 2.0.3, RRID:SCR_012919) (23).

Exploratory analysis of raw read counts was performed using the iDEP (version 1.11) web application (24). Raw counts were filtered with a minimum counts per million (CPM) of 10 in at least two samples; of this, 8275 genes remained. Count data were transformed using EdgeR (version 4.0.15; RRID:SCR_012802) (25) with a pseudo count of four prior to k-means clustering (k=6) of the top 1000 genes (Supplementary Figure 2c) and principal component analysis (PCA). Differential gene expression was performed using DESeq2 (version 1.42.0; RRID:SCR_015687) (26) with an FDR cutoff of 0.05, and a minimum fold-change of 2. Pathway analysis was performed using GAGE (version 2.52.0; RRID:SCR_017067) (27) with minimum geneset size of 30 and an FDR cutoff of 0.05 using the following genesets: Gene Ontology (GO) biological process, GO cellular component, GO molecular function, Reactome, and Kyoto Encyclopedia of Genes and Genomes (KEGG). Significance for pathways was defined as an adjusted *p*-value of < 0.05.

### Quantitative lipidomics

TA muscles from each animal were flash frozen and stored at -80 °C until being shipped on dry ice to Metware Bio (Metware Bio, Woburn, MA, USA) for quantitative lipidomics according to the following procedures. Samples were thawed on ice, and approximately 10 mg of each sample was homogenized using a ball-mill grinder at 30 Hz for 20 s in 1 mL of methyl tert-butyl ether (MTBE):methanol (3:1, v/v) with internal standards and then vortexed for 15 min. The mixture was added to 200 µL of water, vortexed for 1 min, incubated at 4 °C for 10 min, and centrifuged at 13,500 x g for 10 min (4 °C). The upper phase (200 µL) was collected and dried at 20 °C. The residue was reconstituted in 200 µL of acetonitrile:isopropanol (1:1, v/v), vortexed for 3 min, and centrifuged at 13,500 x g for 3 min. The final supernatant (120 µL) was used for the LC-MS/MS analysis.

Ultra-performance liquid chromatography (UPLC) was performed using a Nexera LC-40 (Shimadzu, Kyoto, Japan) with an Accucore C30 (2.6 μm, 2.1 mm × 100 mm) (Thermo Fisher Scientific Inc., Waltham, MA, USA) column at a temperature of 45 °C, flow rate of 0.35 mL/min and injection volume of 2 µL. Linear ion trap (LIT) and triple quadrupole (QQQ) scans were acquired using a triple quadrupole-linear ion trap LC-MS/MS QTRAP 6500+ (Sciex, Concord, ON, Canada) operating in positive and negative ion modes controlled by the Analyst software (Sciex, version 1.6.3; RRID:SCR_015785). All elution gradients presented as percent mobile phase A:percent mobile phase B. Elution gradients: 80:20 v/v at 0 min, 70:30 v/v at 2 min, 40:60 v/v at 4 min, 15:85 v/v at 9 min, 10:90 v/v at 14 min, 5:95 v/v at 15.5 min, 5:95 v/v at 17.3 min, 80:20 v/v at 17.5 min, 80:20 v/v at 20 min.

The electrospray ionization (ESI) source conditions were as follows: source temperature, 500 °C; ion spray voltage (ISV) 5500 V (pos), -4500 V (neg); ion source gas I (GSI) 45 psi; ion source gas II (GSII) 55 psi; curtain gas (CUR) 35 psi. Instrument tuning and mass calibration were performed using 10 and 100 μmol/L polypropylene glycol solutions in the QQQ and LIT modes, respectively. QQQ scans were acquired in multiple reaction mode (MRM) experiments with a collision gas (nitrogen) set to 5 psi. The declustering potential (DP) and collision energy (CE) for individual MRM transitions were determined with further DP and CE optimizations. A specific set of MRM transitions was monitored for each period according to the lipids eluted within this period. A quality control (QC) sample was prepared from a mixture of all sample extracts to examine the reproducibility of the metabolomics process. During data collection, a quality control sample was inserted for approximately every 10 test samples. The percentages of the identified compound classes are shown in Supplementary Figure 3a. The QC data are presented in Supplementary Figure 3b-e.

**Figure 3.**
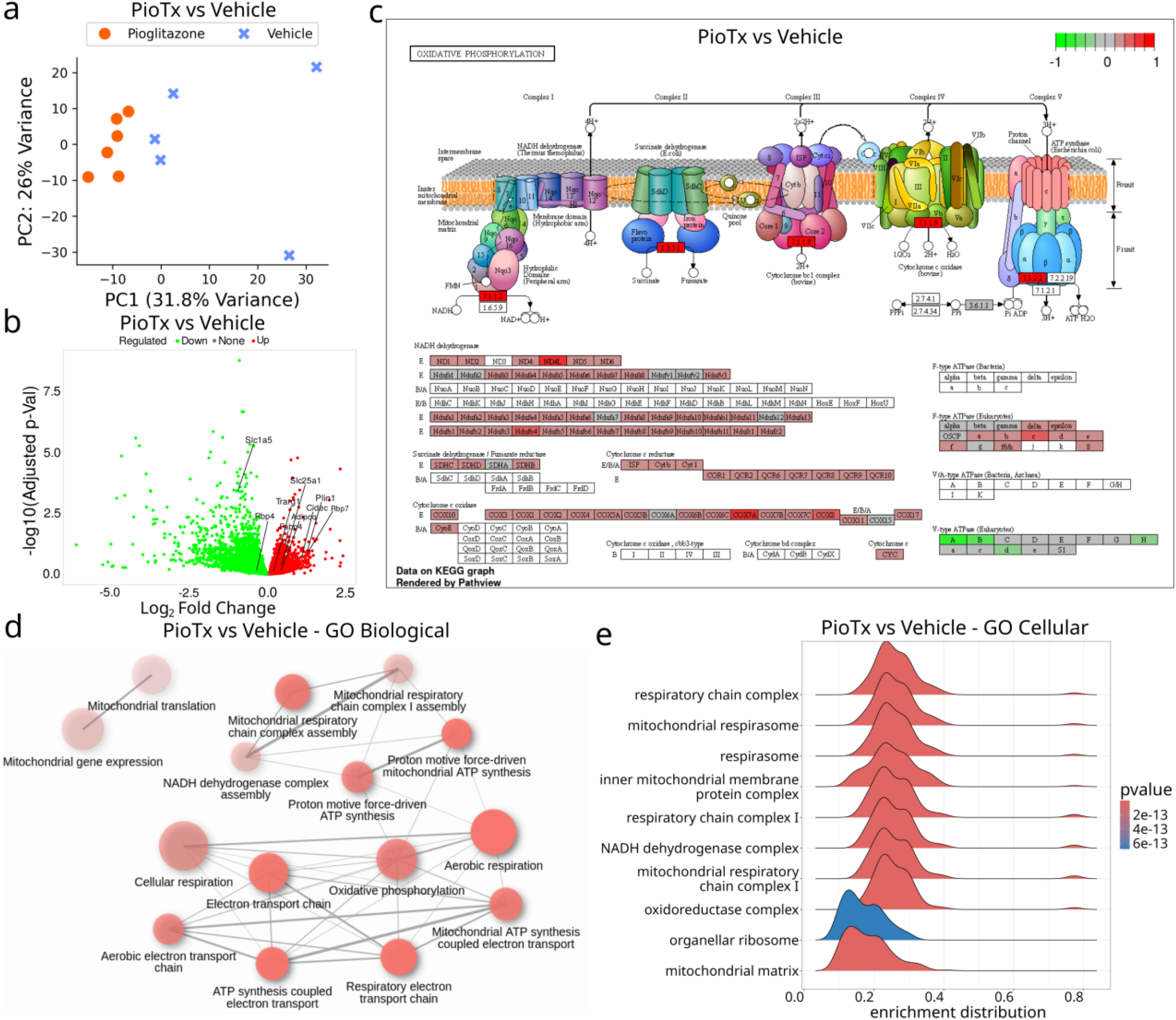
Bulk RNA sequencing of the tibialis anterior. (**a**) 2D PCA plot of bulk RNA-seq from the PioTx (orange, circle; n=6) and vehicle (blue, cross; n=5) groups. (**b**) Volcano plot of downregulated (green, left) and upregulated (red, right) genes comparing the PioTx and vehicle groups. The genes highlighted are downstream target genes of PPARγ. Downregulated: *Slc1a5* and *Rbp4*. Upregulated: *Slc25a1, Cidec, Rbp7, Fabp4, Adipoq, Plin1*, and *Trarg1*. (**c**) KEGG pathway diagram depicting PioTx vs vehicle upregulated (red) and downregulated (green) genes involved in oxidative phosphorylation. (**d**) Network plot of the top 15 GO Biological pathways for PioTx versus vehicle, all of which are upregulated (red). The cutoff threshold for association with another pathway was 30% of genes shared. Opaque nodes represent more significantly enriched gene sets. Thicker connecting lines represent more overlapped genes. (**e**) Ridgeplot of top 10 GSEA enriched GO Cellular pathways for PioTx vs vehicle. NES, normalized enrichment score.

### Lipidomics data analysis

The software Analyst (Sciex, version 1.6.3; RRID:SCR_015785) was used to process raw mass spectral data. Orthogonal partial least squares discriminant analysis (OPLS-DA) models were built for identified metabolites using log2 transform + mean centering for each group comparison using MetaboAnalystR 4.0 (RRID:SCR_016723) (28). To prevent overfitting, 200 permutation tests were performed. The OPLS-DA model validation is shown in Supplementary Figure 4a-c. For the two-group analysis, differential lipids were defined as variable importance in projection (VIP) > 1 and *p*-value (Student’s t-test) < 0.05. All other analyses used unit variance scaling (UV) when applicable with the following equation: x’ = (x-μ)/*σ*. Principal component analysis (PCA) plots are shown in Supplementary Figure 4d-f. The identified lipids were annotated using the KEGG compound database and were mapped to the KEGG pathway database.

**Figure 4.**
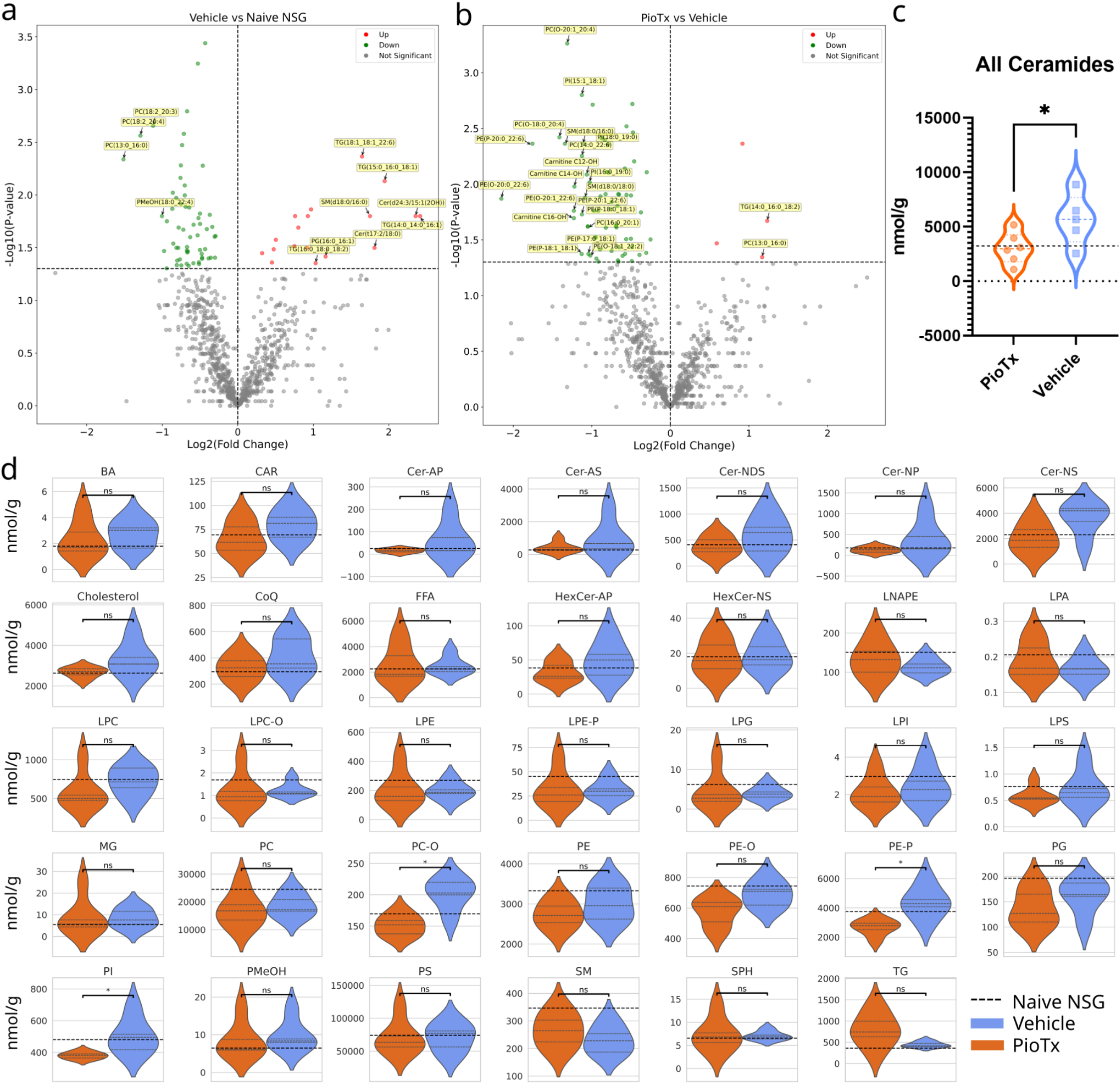
Quantitative lipidomics of the TA muscles. (**a**,**b**) Volcano plots representing log2 fold-change (log2FC) on the horizontal axis and -Log10(*p*-value) on the vertical axis of downregulated (green) and upregulated (red) lipids in vehicle (n=5) vs. naive NSG (n=4) in **a** and PioTx (n=6) vs. vehicle groups in **b**. The horizontal dotted line represents the -log10(*p*-value) equivalent of *p* = 0.05, and the vertical dotted line is at a log2FC of zero. (**c**) Violin plot representing the significant difference in total ceramide abundance between PioTx and vehicle groups. (**d**) Violin plots of raw lipid abundance per subclass in nmol∙g^-1^. (**c**,**d**) The black horizontal dotted line represents the mean lipid abundance for naive NSG. The gray horizontal dotted lines represent quartiles. Complete lipid subclass names are defined in Supplementary Table 5. * < 0.05.

The abundance of lipid subclasses were calculated using summed abundances for all lipids in the subclasses per sample, and then subclass abundance was compared between PioTx and vehicle groups using Student’s t-test. Additional parent classes representing sums of grouped subclasses were also considered. *Total Ceramides* reflect summed abundance of 57 lipids across 7 subclasses (Cer-AP, Cer-AS, Cer-NDS, Cer-NP, Cer-NS, HexCer-AP, HexCer-NS), *Fatty Acids and Derivatives* reflect 124 lipids across 5 subclasses (FFA, MG, TG, LNAPE, LPA), and *Glycerophospholipids* reflect 638 lipids across 16 subclasses (PC, PC-O, PE, PE-O, PE-P, PG, PI, PS, LPC, LPC-O, LPE, LPC-O, LPE, LPE-P, LPG, LPI, LPS, PMeOH).

### Untargeted and widely targeted metabolomics

The same TA muscles used in quantitative lipidomics were flash frozen and stored at -80 °C until being shipped on dry ice to Metware Bio for untargeted and widely targeted metabolomics according to the following procedures. Samples were thawed on ice, and approximately 20 mg of each sample was homogenized using a ball-mill grinder at 30 Hz for 20 s in 400 µL methanol:water (7:3, v/v) with internal standards added to the ground sample and mixed by shaking at 2500 rpm for 5 min. After 15 min on ice, samples were centrifuged at 13,500 x g for 10 min (4 °C), and 300 µL of supernatant was collected and stored at -20 °C for 30 min. The samples were centrifuged again at 13,500 x g for 3 min (4 °C) and a 200 µL aliquot was used for LC-MS analysis.

For untargeted metabolomics, UPLC was performed using an ExionLC 2.0 (Sciex) with an ACQUITY HSS T3 (2.1 mm × 100 mm, 1.8 µm) (Waters, Milford, MA, USA) column at a temperature of 40 °C, flow rate of 0.4 mL/min and injection volume of 5 µL. The settings were the same for widely targeted metabolomics, except for an injection volume of 2 µL. Mass spectrometry was performed using a Quadrupole-Time of Flight TripleTOF 6600+ (Sciex) for untargeted and a tandem mass spectrometer (MS/MS) QTRAP® 6500+ (Sciex, RRID:SCR_021831) for widely targeted metabolomics, both operating in positive and negative ion mode controlled by Analyst software (Sciex; version 1.6.3). All elution gradients are presented as percent mobile phase A:percent mobile phase B. Untargeted elution gradients: 95:5 v/v at 0 min, 10:90 v/v at 11 min, 10:90 v/v at 12 min, 95:5 v/v at 12.1 min, and 95:5 v/v at 14 min. Widely-targeted elution gradients: 95:5 v/v at 0 min, 10:90 v/v at 10 min, 10:90 v/v at 11 min, 95:5 v/v at 11.1 min, 95:5 v/v at 14 min. For both untargeted and widely targeted methods, mobile phase A was ultrapure water with 0.1% formic acid, and mobile phase B was acetonitrile with 0.1% formic acid.

The untargeted electrospray ionization (ESI) source conditions were as follows: source temperature, 500 °C; ISV 5500 V (pos), -4500 V (neg); GSI 50 (pos/neg); GSII 50 (pos/neg); CUR 25 (pos/neg); DP 80 (pos), -80 (neg); CE 30 (pos), -30 (neg); and collision energy speed (CES) 15 (pos/neg). The widely targeted ESI conditions: source temperature, 500 °C; ISV 5500 V (pos), -4500 V (neg); GSI 50 psi; GSII 50 psi; CUR 25 psi; and collision gas (CAD) was high. Instrument tuning and mass calibration were performed using 10 and 100 μmol/L polypropylene glycol solutions in triple-quadrupole (QQQ) and linear ion trap (LIT) modes, respectively.

A specific set of MRM transitions was monitored for each period according to the metabolites eluted within this period. Similar to quantitative lipidomics, the percentages of the identified compound classes are shown in Supplementary Figure 5a, and QC samples were prepared in the same manner and the data are presented in Supplementary Figure 5b-e.

**Figure 5.**
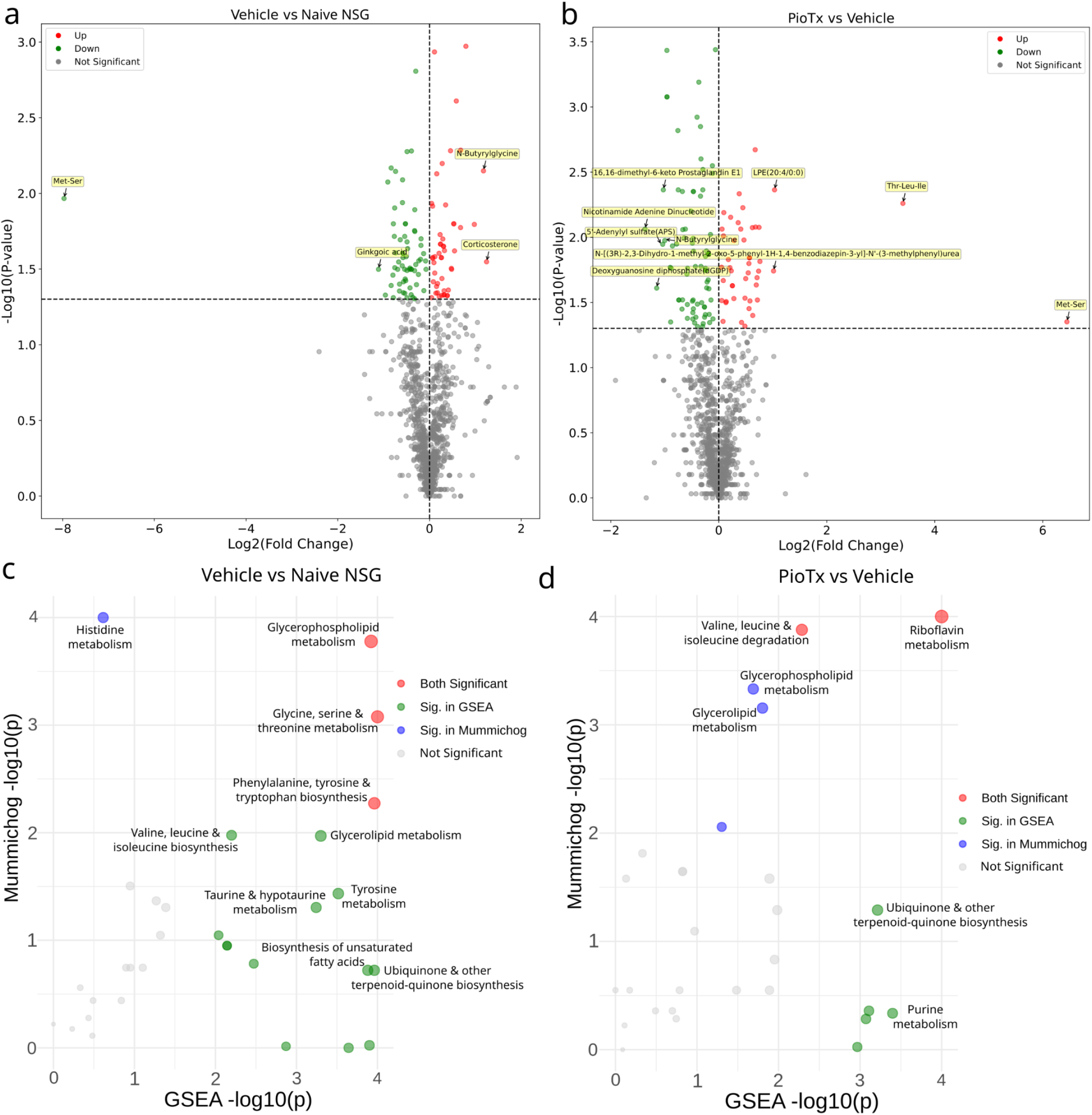
Untargeted metabolomics of the TA muscles. (**a**,**b**) Volcano plots representing log2 fold-change (log2FC) on the horizontal axis and -Log10(*p*-value) on the vertical axis of downregulated (green) and upregulated (red) metabolites in vehicle (n=5) vs. naive NSG (n=4) in **a** and PioTx (n=6) vs. vehicle in **b**. The horizontal dotted line represents the -log10(*p*-value) equivalent of *p* = 0.05, and the vertical dotted line is at a log2FC of zero. (**c**) Scatter plot of significant integrated pathway activity results from both GSEA (horizontal axis) and mummichog (vertical axis) in vehicle versus naive NSG. *P*-values are shown as -log10 transformed values. (**d**) Scatter plot of significant integrated pathway activity results from both GSEA (horizontal axis) and mummichog (vertical axis) in PioTx versus vehicle. *P*-values are shown as -log10 transformed values. (**c**,**d**) MetaboAnalyst 6.0 was used for pathway analysis and figure generation. Bottom-left quadrant represents not significant pathways, top-left (blue circles) significant mummichog pathways, bottom-right (green circles) significant GSEA pathways, and top-right (red circles) significant pathways in both GSEA and mummichog. The significance threshold for both combined pathway analyses was ɑ = 0.05.

### Metabolomics data analysis

Raw peak intensities (intensity unit: CPS, counts per second) were preprocessed by imputing missing values using ⅕ th of the minimum value of each metabolite. The coefficient of variation (CV) of the quality control (QC) sample was calculated, and metabolites with CV < 0.3 were retained as final metabolites. Analyst software (Sciex, version 1.6.3; RRID:SCR_015785) was used to process the raw mass spectral data. Internal standards (IS) were added to QC samples, stability was assessed, and the CV of all IS was < 15%, with the highest value being 2.3% (Supplementary Data 1). The OPLS-DA models were built as previously described, and the model validation is shown in Supplementary Figures 6a-c. For the two-group analysis, differential metabolites were defined as variable importance in projection (VIP) > 1 and *p*-value (Student’s t-test) < 0.05. All other analyses used unit variance scaling (UV) when applicable with the following equation: x’ = (x-μ)/*σ*. PCA plots are shown in Supplementary Figure 6d-f. The identified metabolites were annotated using the KEGG compound database and were mapped to the KEGG pathway database. Pathways containing significantly dysregulated metabolites were analyzed using metabolite set enrichment analysis (MSEA), and significance was determined using hypergeometric tests. Combined pathway analysis was performed using Mummichog and GSEA in MetaboAnalyst 6.0 (RRID:SCR_015539) with a pathway significance cutoff of 0.05.

**Figure 6.**
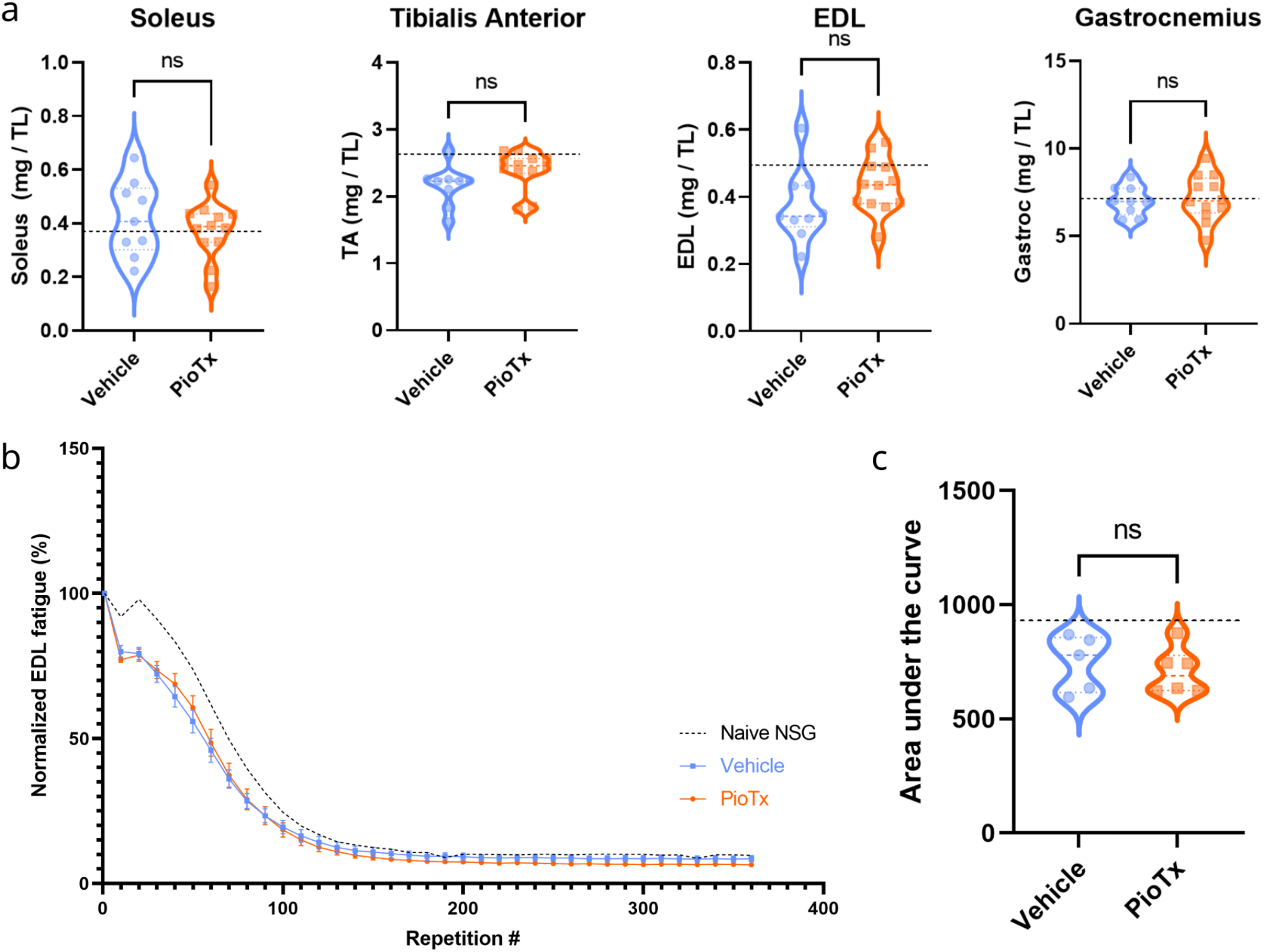
*Ex vivo* muscle weight and EDL functional testing. (**a**) Skeletal muscle weights (mg) for vehicle (blue, left; n=9) and PioTx groups (orange, right; n=12) normalized to tibial length (mm) from left to right for the soleus, tibialis anterior (TA), extensor digitorum longus (EDL), and gastrocnemius, respectively. No significant differences were observed between the groups in any muscle. Black dotted lines represent the mean normalized muscle weights of the soleus (0.38), TA (2.61), EDL (0.49), and gastrocnemius (7.1) for a naive NSG mouse. (**b**) Normalized EDL fatigue curves for PioTx (n=6) and vehicle (n=5) groups. Two-way repeated-measures ANOVA of EDL contractions normalized to the first repetition. Error bars represent standard error of the mean (SE). (**c**) Area under the curve (AUC) for data presented in **b** for the vehicle (blue, left) and PioTx (orange, right) groups. There were no significant differences between the groups. (**b**,**c**) Black dotted lines represent the mean EDL fatigue and corresponding mean AUC (mean=918.10) for a naive NSG mouse. TL, tibia length.

### Whole-animal indirect calorimetry and metabolic monitoring

To evaluate the live whole-animal metabolic activity, tumor groups were housed for 5 days in the Oxymax-CLAMS system (Comprehensive Laboratory Animal Monitoring System; Columbus Instruments, Columbus, OH, USA; RRID:SCR_016718). The environment was maintained at 22 °C with a sampling flow rate of 0.5 liters per minute (LPM) and 800.98 mmHg of pressure, and the system was calibrated prior to each experimental run. CLAMS was performed during the third week of pioglitazone treatment, which was provided in drinking water at the same dosage as the oral gavage (30 mg·kg^-1^·day^-1^). Following a 13-17 hour acclimation period, animals were monitored for 96 consecutive hours. One PioTx mouse and one vehicle mouse were excluded from analysis for data integrity concerns arising from sampling discrepancies. Raw Oxymax-CLAMS data were pre-processed for downstream statistical analysis using CLAMS Wrangler (version 1.0.5) and CalR (version 1.3) (29) (30). The primary variables of interest were mean VO_2_ (ml∙hr^-1^), VCO_2_ (ml∙hr^-1^), respiratory exchange ratio (RER), cumulative X- and Y-axis infrared beam breaks, cumulative food consumption, energy expenditure (EE), and energy balance. Animal weights (grams) at the time of entry into Oxymax-CLAMS, and a food energy density of 3.1 kcal∙g^-1^ was used for calculating EE and energy balance.

### *Ex vivo* muscle function testing

Our mouse model of BC induced fatigue has been previously established (14) (31). Anesthesia was induced with 4% isoflurane and maintained at 2.5%. The following muscles were bilaterally dissected for the evaluation of absolute mass: TA, extensor digitorum longus (EDL), gastrocnemius, and soleus. Tibia length was measured via calipers for muscle weight normalization. *Ex vivo* isometric analysis was performed on the EDL and soleus using established laboratory protocols (32) (33). In brief, muscles were immediately transferred to an oxygenated muscle stimulation bath containing Ringer’s solution (100 mM NaCl, 4.7 mM KCl, 3.4 mM CaCl_2_, 1.2 mM KH_2_PO_4_, 1.2 mM MgSO_4_, 25 mM HEPES, and 5.5 mM D-glucose) that was maintained at 4 °C. Muscle function testing was performed at 22 °C using the Aurora Scientific 1300A 3-in-1 Whole Animal System (mouse) running Aurora Scientific 605A Dynamic Muscle Data Acquisition software (Aurora Scientific, Aurora, ON, Canada). The muscle length was gradually increased to obtain the maximal twitch force response; this muscle length was recorded as the optimal length (L_o_). Muscle cross-sectional area (CSA) was calculated by dividing the muscle mass by the product of the muscle density coefficient (1.06 g·cm^3^), muscle L_o_, and the fiber length coefficient (EDL: 0.45, soleus: 0.69), this CSA value was used to calculate muscle-specific force (i.e., force mN·muscle CSA^−1^) (34) (35). Muscle contractile parameters obtained from isometric twitch contractions included the peak isometric twitch force, contraction time (CT), ½ relaxation time (½ RT), rate of force development (RFD), and rate of relaxation (RR).

With the muscle set at L_o_, muscles were stimulated with 500 ms tetanic trains at increasing frequencies (5, 10, 25, 50, 80, 100, 120, and 150 Hz) to establish the force–frequency relationship. Each contraction was followed by 2 min rest. Absolute isometric tetanic force was recorded at each stimulation frequency. Assessment of skeletal muscle fatigue was performed using repeated 40 Hz tetanic trains that occurred once per second lasting 330 ms each, for a total of 360 repetitions across 6 minutes (32). Peak force generated per repetition was normalized to the first repetition force and reported as percent (%) change in force. With normalized peak force plotted for each repetition, the area under the fatigue curve (AUC) was calculated for each muscle.

### Statistical analyses

GraphPad Prism (version 10; RRID:SCR_002798) was used for statistical analyses, with significance set at an alpha value of 0.05, unless otherwise specified. Two-tailed unpaired Student’s t-tests were used to compare muscle weights normalized to tibia length, AUC, final tumor volumes, final body weights, and EDL and soleus muscle isometrics, as described in Supplementary Tables 2 and 3, respectively. EDL and soleus fatigue were analyzed using a repeated-measures two-way ANOVA with Geisser-Greenhouse correction and Šidák multiple comparisons test. Tumor growth kinetics were analyzed using the exponential regression model V=V_0_e^kt^ where V represents volume in mm^3^, V_0_ is the volume at the start of treatment in mm^3^, k is the rate constant in days^-1^, and t represents time in days relative to the start of treatment. Muscle force-frequency relationships were analyzed using the 4 parameter logistic regression model P = P_min_ + ((P_max_ - P_min_)/(1 + (K_f_/f)^h^)), where P represents muscle force in mN and f represents stimulation frequency in Hz. The following parameters were obtained from the force–frequency curve: minimum force (P_min_), maximum force (P_max_), half-frequency (K_f_), and the Hill coefficient (h). K_f_ is defined as the frequency at which the developed force is the midpoint between P_min_ and P_max_, where h describes the slope of the force–frequency logistic curve <u>(36)</u>. Regression parameters were compared between groups using two-tailed unpaired Student’s t-tests. Two-tailed Student’s t-tests were performed on CLAMS metabolic variables to evaluate differences between vehicle and PioTx groups across the full day period.

## Results

### Study overview and characterization of BC-PDOX mouse model

To validate equal tumor burden between the groups, we performed weekly ultrasound measurements and body weight assessments. Figure 1a illustrates the experimental timeline, while Figure 1b outlines the utilization of isolated muscles. For a consistent analysis, bulk RNA sequencing and metabolomics/lipidomics assays were conducted on the contralateral tibialis anterior (TA) muscles of each animal. The extensor digitorum longus (EDL) and soleus muscles, representing fast (Type II) and slow (Type I) twitch muscles, respectively, were chosen for functional testing because of their differing fatigue responses. Tumor growth kinetics did not differ between groups (*p* = 0.0870) with a V_0_ of 728 mm^3^ (SE ± 21.9) and a doubling time of 22.0 days (SE ± 1.01) for the BC-PDOX Captisol-treated (vehicle) group (n=5) and a V_0_ of 705 mm^3^ (SE ± 19.5) and a doubling time of 23.7 days (SE ± 1.05) for the BC-PDOX pioglitazone-treated (PioTx) group (n=6) (Fig. 1c). Terminal tumor volumes were also statistically equivalent between groups (Student’s t-test, *p* = 0.503), averaging 1647 mm^3^ (SD ± 258.2) for the vehicle group and 1519 mm^3^ (SD ± 335.0) for the PioTx group. Similarly, body weight did not differ (*p* = 0.497), with the vehicle group averaging 25.9 g (SD ± 1.6) and the PioTx group averaging 26.6 g (SD ± 1.5) (Supplementary Data 2). The original and passaged tumors demonstrated over 80% concordance in their short tandem repeat (STR) profiles, suggesting the stability of the tumor model in the fifth implanted passage (Supplementary Figure 1). NSG-specific genotyping probes (Il2rg WT, Il2rg KO, Scid Mutation) confirmed the strain identity of all the mice. These results validated the BC-PDOX model, as evidenced by consistent tumor burdens and body weights across groups, as well as no large differences in the expected tumor or mouse strain identity.

### Whole-animal indirect calorimetry

Whole-animal indirect calorimetry was performed using the Oxymax-CLAMS system to examine the systemic metabolic effects of the pioglitazone treatment. Body mass at entry into Oxymax-CLAMS did not differ between treatment groups (*p* = 0.988), nor did the volume of water consumed while in Oxymax-CLAMS (*p* = 0.127). A significantly lower average O_2_ consumption (ml∙hr^-1^) (*p* = 0.035) (Fig. 2a), and CO_2_ production (ml∙hr^-1^) (*p* = 0.022) (Fig. 2b) was observed in the PioTx mice. Similarly, cumulative energy expenditure (EE) (kcal∙hr^-1^) was significantly lower in PioTx mice across the four day period (*p* = 0.035) (Fig. 2c), however, total food consumed throughout this time did not differ (*p* = 0.788) (Fig. 2d). As such, cumulative energy balance across the four day period was significantly lower in the vehicle group (*p* = 0.014), reflecting a larger net energy deficit in vehicle mice. In addition to the presented analysis, there was no significant difference between groups for respiratory exchange ratio (RER) (*p* = 0.860), locomotor activity (*p* = 0.998), or ambulatory activity (*p* = 0.670).

### Bulk RNA-seq of TA muscles

Bulk RNA sequencing was performed to confirm the expected action of pioglitazone interacting with PPARγ and its subsequent influence on downstream gene targets. Principal component analysis (PCA) revealed separation between the PioTx and vehicle groups, indicating distinct gene expression profiles. Principal component (PC) 1 explained 31.8% of the variation and PC2 explained 26% of the variation (Fig. 3a; Supplementary Figure 2d). Unnormalized gene expression counts are provided in Supplementary Data 3.

#### PPAR target genes

K-means cluster 5 showed enrichment of the PPAR signaling pathway (*adj. p* < 0.05, fold change 6, KEGG:mmu03320), which was expected as a result of pioglitazone treatment (Supplementary Figure 2e). Cluster 5 included several dysregulated genes within the PPAR pathway, including *Fabp4, Fabp3, Pltp, Scd1, Scd2*, and *Aqp7* (Supplementary Data 4). Regulation of established downstream genes of PPARγ: *Slc1a5* and *Rbp4* appear downregulated following pioglitazone-treatment, whereas *Slc25a1, Cidec, Rbp7, Fabp4, Adipoq, Plin1*, and *Trarg1* appear upregulated (Fig. 3b). Gene set enrichment analysis (GSEA) further supported the enrichment of the PPAR signaling pathway, with a normalized enrichment score (NES) of 1.37; however, the FDR q-value was not significant at 0.071 (Supplementary Figure 2f).

#### Genes involved in oxidative phosphorylation

Kyoto Encyclopedia of Genes and Genomes (KEGG) sources showed upregulation of the oxidative phosphorylation (KEGG:mmu00190) pathway in PioTx compared to vehicle (*adj. p* < 0.001) (Fig. 3c; Supplementary Table 4). In particular, the vast majority of genes involved in the electron transport chain (ETC) and ATP synthase were upregulated to some extent, with the most upregulated being *Nd4l* and *Cox7a* (Fig. 3c). Other relevant upregulated pathways included citrate cycle (TCA cycle) (KEGG:mmu00020), carbon metabolism (KEGG:mmu01200) and fatty acid metabolism (KEGG:mmu01212) (all *adj. p* < 0.05) (Supplementary Data 5).

#### Mitochondrial bioenergetic pathways

PioTx mice showed upregulation of mitochondrial bioenergetic-related pathways compared with vehicle mice. Pathway analysis was performed using multiple gene set sources: Gene Ontology (GO) Biological (Bio), Cellular (Cell), and Molecular (Mol); Reactome; and KEGG. Significantly upregulated GO Bio pathways related to mitochondrial bioenergetics included, but were not limited to: proton motive force-driven mitochondrial ATP synthesis (GO:0042776), ATP synthesis coupled electron transport (GO:0042773), respiratory electron transport chain (GO:0022904), and aerobic respiration (GO:0009060) (all *adj. p* < 0.001) (Fig. 3d; Supplementary Table 4; Supplementary Data 6). Similar trends were observed in PioTx GO Cell pathways. Significantly upregulated pathways included, but were not limited to: oxidoreductase complex (GO:1990204), inner mitochondrial membrane protein complex (GO:0098800), mitochondrial respirasome (GO:0005746), and NADH dehydrogenase complex (GO:0030964) (all *adj. p* < 0.001) (Fig. 3e; Supplementary Table 4; Supplementary Data 7). Pathways from Reactome further supported the overall observed trend of mitochondrial bioenergetic pathway upregulation following pioglitazone treatment, including complex I biogenesis (R-MMU-6799198), respiratory electron transport (R-MMU-611105), and pyruvate metabolism and Citric Acid TCA cycle (R-MMU-71406) (all *adj. p* < 0.001) (Supplementary Table 4; Supplementary Data 9). All adjusted *p*-value significant pathways for each genset source are shown in Supplementary Data 5-9. Overall, pioglitazone administration was associated with the upregulation of numerous pathways involved in mitochondrial bioenergetics, as corroborated by multiple gene set analyses.

### Quantitative lipidomics in tibialis anterior

Quantitative lipidomics (QL) was performed to explore changes in lipid abundance. A total of 901 lipids were detected across all samples after data filtering. Differential lipids were as follows: PioTx vs. vehicle, 66 down and 4 up; vehicle vs. naive NSG, 60 down and 18 up; PioTx vs. naive NSG, 116 down and 9 up. The total abundance of lipid content showed slight variation between groups, with the naive NSG having the highest abundance (117,734 nmol∙g^-1^), followed by the vehicle (111,762 nmol∙g^-1^) and PioTx (105,282 nmol∙g^-1^).

Differential lipids in vehicle vs. naive NSG and PioTx vs. vehicle groups are shown in Fig. 4a and 4b, respectively. Vehicle vs. naive NSG shared 7 of the significantly dysregulated lipids with PioTx vs. vehicle. Of the top 20 differential lipids in the vehicle vs. naive NSG group, 11 were upregulated and 9 were downregulated. In PioTx vs. vehicle, 2 were upregulated and 18 were downregulated.

#### Ceramides

One of the most prominent differences observed in lipid abundance is that of the ceramides, which in our dataset is composed of the classes ceramide (Cer) and hexosylceramide (HexCer). The overall abundance of *Total Ceramides* (Cer+HexCer) was significantly lower in the PioTx group compared to the vehicle (*p* = 0.048) (Fig. 4c). Ceramide subclasses include alpha-hydroxy fatty acid-phytosphingosine (AP) (*p* = 0.173), alpha-hydroxy fatty acid-sphingosine (AS) (*p* = 0.272), non-hydroxy fatty acid-dihydrosphingosine (NDS) (*p* = 0.185), non-hydroxy fatty acid-phytosphingosine (NP) (*p* = 0.167), and non-hydroxy fatty acid-sphingosine (NS) (*p* = 0.074). HexCer subclasses include HexCer-AP (*p* = 0.173) and HexCer-NS (*p* = 0.848). The abundance for the 7 individual ceramide subclasses did not significantly differ, though some trended towards a greater abundance in the vehicle group (Fig. 4d).

*Other subclass abundances*. The differences in lipid abundance between the subclasses across the sample groups are plotted in Fig. 4d. Parent groups *Fatty Acids and Derivatives* (*p* = 0.516) and *Glycerophospholipids* (*p* = 0.814) did not differ between PioTx and vehicle groups. Individual subclasses that were significantly different between groups were all within the Glycerophospholipid parent class, including alkyl-phosphatidylcholine (PC-O) (*p* = 0.003), alkenyl-phosphatidylethanolamine (PE-P) (*p* = 0.007), and phosphatidylinositol (PI) (*p* = 0.025), in which the vehicle group was significantly greater in abundance. Other subclasses that trended towards significance include greater abundances in the vehicle group for cholesterol (*p* = 0.095) and alkyl-phosphatidylethanolamine (PE-O) (*p* = 0.056), and a greater abundance in the PioTx group in triacylglycerol (TG) (*p* = 0.083). All other subclasses did not differ in lipid abundance between PioTx and vehicle groups.

### Untargeted metabolomics in tibialis anterior

To explore the changes in metabolite abundance as a result of pioglitazone treatment, we performed untargeted metabolomics. Across all samples, we detected 1186 metabolites after data filtering. Differential metabolites were as follows: PioTx vs. vehicle, 79 down and 46 up; vehicle vs. naive NSG, 65 down and 49 up; PioTx vs. naive NSG, 133 down and 73 up. Vehicle vs. naive NSG shared 14 of the significantly dysregulated metabolites with PioTx vs. vehicle.

#### Differential metabolites

Differential metabolites in vehicle vs. naive NSG and PioTx vs. vehicle groups are shown in Fig. 5a and 5b, respectively. Comparison of the top 10 differential metabolites between PioTx vs. vehicle and vehicle vs. naive NSG revealed two common metabolites, N-butyrylglycine and Met-Ser (methionyl-serine). N-Butyrylglycine was significantly upregulated in vehicle vs. naive NSG (*p* = 0.00710, Log2FC = 1.18) and downregulated in PioTx vs. vehicle (*p* = 0.0106, Log2FC = -1). Met-Ser is significantly downregulated in vehicle vs. naive NSG (*p* = 0.0108, Log2FC = -7.97) and proportionally upregulated following pioglitazone (*p* = 0.0446, Log2FC = 6.45). The reversal of these metabolite levels suggests a potential role of pioglitazone in modifying the metabolic state of SkM affected by BC.

#### Amino acid metabolism

Combined pathway analysis using mummichog and GSEA identified numerous dysregulated pathways involved in amino acid metabolism when comparing vehicle vs. naive NSG (Fig. 5c). Metabolite set enrichment analysis (MSEA) comparing vehicle and naive NSG further supports amino acid metabolism disruption (Supplementary Figure 7a). However, pioglitazone did not appear to ameliorate dysregulation of these pathways (Fig. 5d, Supplementary Figure 7b).

### Muscle fatigue and isometric analysis

Muscle function testing was performed *ex vivo* to quantitatively evaluate isometric and fatigue properties in the EDL and soleus, which represent fast-twitch and slow-twitch predominant muscles, respectively.

#### Muscle weights

All muscles were weighed immediately after isolation. Muscle weights were normalized to tibial length to account for variations in body size. No significant differences in normalized weights between the PioTx and vehicle groups were found in the EDL (*p* = 0.147), soleus (*p* = 0.416), TA (*p* = 0.083), and gastrocnemius (*p* = 0.741) (Fig. 6a, Supplementary Table 6).

#### Fatigue testing

Figure 6b shows the normalized EDL force output (percent change from the first rep) over the course of 360 isometric contractions occurring at 40 Hz. The fatigue curves for each group showed no significant separation. No significant difference in the AUC was found between the PioTx and vehicle groups (*p* = 0.596) (Fig. 6c). The same analysis revealed no shifts in the fatigue curve for the soleus muscle or significant differences in the AUC between groups (Supplementary Figure 8a-b).

#### Isometric contractile metrics

Pioglitazone treatment did not result in any significant differences in any of the measured EDL or soleus isometric contractile properties (Supplementary Tables 2 and 3, respectively). The force-frequency relationship (FFR), which represents the interaction between muscle force production and the stimulation frequency used to induce muscle contraction, was established for the fast EDL and slow soleus muscles. We observed no differences in the overall FFR in the fast EDL and slow soleus muscles in our study when comparing PioTx and vehicle groups (Supplementary Figure 8c-f).

## Discussion

In this preclinical drug trial, we examined the metabolic effects of pioglitazone treatment in SkM derived from PDX-implanted mice as a potential therapeutic agent for the treatment of BC-induced fatigue. Whole-animal indirect calorimetry showed significant decreases in O_2_ consumption, CO_2_ production, and energy expenditure after pioglitazone treatment, which is consistent with the known effects of pioglitazone on metabolism (37) (38). Overall, these findings suggested a decrease in metabolic activity as a result of pioglitazone treatment.

Additionally, RER showed no significant differences between the groups, indicating no shift in substrate utilization for energy production. Based on our data, decreased metabolic rate was not correlated with less activity.

Beyond improving insulin resistance, pioglitazone has been shown to improve skeletal muscle mitochondrial function in type 2 diabetic (T2D) mice by increasing ADP-dependent mitochondrial respiration, complex I and III activities, and reducing oxidative stress (39). In human skeletal muscle from T2D patients, pioglitazone improves fatty acid metabolism and phosphocreatine usage (40), along with increasing the abundance of some ATP synthesis-related proteins (41). Research into the effects of pioglitazone in treating BC-induced SkM alterations is limited, but our laboratory has previously shown that as little as 2 weeks of pioglitazone therapy is associated with the restoration of mitochondrial-associated pathways in skeletal muscles of BC-PDOX mice at the transcript level (15).

Following 4-weeks of pioglitazone treatment, there was considerable restoration in the transcriptomic profile of the SkM from PioTx mice despite the presence of BC without an observed concomitant restoration in fatigue via *ex vivo* testing. This lack of improvement suggests that permanent or long-term changes may contribute to fatigue. One possible reason for this is alterations in the availability, structure, or function of proteins involved in biogenesis. While we demonstrated gene upregulation, the turnover rate for proteins in mouse cardiac mitochondria can range from hours to months (42), suggesting that any defects in mitochondrial protein function may persist beyond the 28 day treatment duration. In addition to changes in bioenergetic pathways, alterations in structural and contractile proteins within skeletal muscle may also be a contributing factor that warrants further study.

Dysregulation of lipid subclasses has been previously associated with cancer cachexia. One study examined plasma from both cachectic mice and humans and found decreases in LPC lipid species and increases in ceramides (43). Our data in skeletal muscles followed the same trend of elevated ceramides when comparing vehicle to naive NSG mice with overall total ceramide abundance being increased in the vehicle group.

Pioglitazone treatment significantly decreased total ceramide levels compared to vehicle treated mice. One role of ceramides is to serve as a secondary messenger in the sphingomyelin (SM) signaling pathway. Despite the significant decrease in total ceramide abundance following pioglitazone treatment, PioTx SM levels mirror the vehicle group, both of which are lower than naive NSG. Interestingly, the literature shows a relationship between insulin resistance and prominent ceramide accumulation (44), although one study showed higher levels in predominantly oxidative muscles and minimal changes in glycolytic (45), whereas our study examined a predominantly glycolytic muscle. Intramyocellular accumulation of ceramides has been found to disrupt insulin signaling (46). The marked increase in ceramide levels seen in the tumor-bearing vehicle-treated mice should be explored in future studies, as well as the potential contributions to fatigue.

N-Butyrylglycine was found to be upregulated in vehicle mice compared to naive NSG and was subsequently downregulated following pioglitazone treatment. It is a metabolite of fatty acid breakdown that can be found in body fluids, the elevation of which is associated with mitochondrial fatty acid β-oxidation (FAO) dysfunction, particularly short-chain acyl-CoA dehydrogenase deficiency (SCAD) (47). The amino acid intermediate metabolite methionyl-serine (Met-Ser) was greatly downregulated in the vehicle group and proportionally upregulated following pioglitazone treatment. Although the literature directly mentioning Met-Ser is sparse, dysfunction of amino acid metabolism was a common theme in our data. MSEA revealed three significantly enriched pathways in vehicle vs. naive NSG: pantothenate and CoA biosynthesis; valine, leucine, and isoleucine degradation; and valine, leucine, and isoleucine biosynthesis (all FDR-corrected *p* < 0.05). The hit compounds in the pantothenate and CoA biosynthesis pathways were L-Valine, pantetheine, and uracil.

However, these pathways were not significantly enriched in PioTx vs. vehicle, suggesting that pioglitazone treatment has minimal restorative benefits on these pathways.

Given the upregulation of a significant number of mitochondrial bioenergetic-associated pathways following pioglitazone therapy, it was disappointing that we were unable to detect preservation of muscle force production in our fatigue protocol following pioglitazone treatment in BC-PDOX mice. The fatigue protocol was based on the seminal paper by Burke et. al. (48), in which motor units were stimulated at 40Hz for 330ms every second. These stimulation parameters were specifically selected to differentiate between the effects of force output due to repeated neuromuscular activation and failure of fiber activation. Our data in isolated EDL skeletal muscles are consistent with this original report in that force decline was observed after 30 s and was dramatically lower after 60-90 s of stimulation (48). When we analyzed the FFR, a 40 Hz stimulation of an isolated EDL muscle would produce approximately 35-40% of the maximal tetanic force. Thus, each stimulation of the fatigue protocol is a submaximal contraction of less than 50% of the theoretical maximum force. Although Burke was able to use this protocol to characterize fatigability differences in individual motor units (48), the protocol may not be sensitive enough to determine improvements in force production due to any specific energy system contribution. The fatigue protocol is likely affected by the interplay of the three main energy systems in skeletal muscle: ATP-PCr, Glycolysis, and Oxidative Phosphorylation. It has been estimated that energy supply to a working muscle during a 30 s sprint can be accounted for by 53% ATP-PCr, 44% glycolysis and 28% mitochondrial respiration (49) (50). Therefore, we suggest that although we have been able to use this protocol to quantify a greater rate of force loss following tumor growth (51) (14), this protocol is likely affected by numerous interconnected mechanisms within skeletal muscle that underlie metabolism and energy production. Furthermore, while mitochondrial respiration likely contributes to energy production during this fatigue protocol, the relative contribution of these pathways during the first 1-2 minutes of the protocol may be too low to be quantified. We are actively investigating additional methods for assessing muscle fatigability in our BC-PDOX mouse model.

## Limitations

Although this study provides novel insights into the effects of pioglitazone treatment on BC-induced changes in the SkM metabolome and lipidome, there are some limitations that should be considered. Due to inherent variations in the rate of tumor progression, animals did not start and end whole-animal indirect calorimetry uniformly. While housed in the Oxymax-CLAMS system, pioglitazone was delivered in drinking water and not by oral gavage, so the dosage consumed varied slightly during this period. After conversion from human to animal dosage, as described here (52), the dosage of pioglitazone administered is approximately five times that approved for humans, which may limit translatability. We only examined one tumor subtype (HER2/neu overexpressing) in this study, and thus could not speak to other similar or differing effects in other BC subtypes. The necessity of using an immunodeficient mouse to successfully engraft PDX tumors precludes any insight into the influence of the immune system on the -omics examined in our model.

## Future directions

PPARγ expression within SkM is relatively low. Our laboratory has demonstrated that IL-15 treatment of muscle in vitro induces the expression of PPARδ and PGC1α in a dose- and time-dependent manner (53), as well as PPARγ and PPARα (unpublished observations). Transgenic mice with greater IL-15 content in SkM also contained correspondingly greater PPARδ and PGC1α (32). With IL-15 in development as an immuno-oncology agent (54) (55), we are pursuing the concept of IL-15 pre-treatment for the induction of PPAR expression in SkM to potentiate the effects of pioglitazone. Other TZDs have a higher binding affinity for PPARγ than pioglitazone, such as rosiglitazone and lobeglitazone, the latter of which is 12x greater resulting in a lower effective dose (56). However, the lack of FDA approval or market acceptance limits the translatability of trialing these drugs. The field would benefit from future studies exploring the use of TZDs or other drugs and concomitant chemotherapy to explore the reduction in fatigue potential while undergoing active treatment. A logical next step would be to further explore the effects of BC on SkM metabolism to elucidate the full extent of dysregulation. This should be done using multiple BC subtypes and immunocompetent models to understand the contribution of the immune system to metabolic dysregulation in SkM. The secondary interaction of pioglitazone with the outer mitochondrial membrane protein, mitoNEET, and its potential role in the metabolic changes presented here should be explored further.

## Conclusions

In summary, this study found that 4 weeks of pioglitazone treatment in HER2/neu overexpressing patient-derived xenograft implanted mice resulted in decreased metabolic activity, upregulation of mitochondrial bioenergetic pathways, partial restoration of dysregulated metabolites and lipid content, and no functional improvement of *ex vivo* muscle fatigue. Further research is needed to fully elucidate the extent of metabolic and lipid dysregulation in skeletal muscle resulting from BC. Pioglitazone may play a role in the successful treatment of BC-induced skeletal muscle dysregulation that leads to fatigue. Future studies should focus on profiling systemic metabolic dysfunction, identifying molecular biomarkers of fatigue, and testing alternative pioglitazone treatment regimens.

## Supporting information

Supplemental Data List

## Acknowledgments

We are deeply grateful to the breast cancer patients who generously provided their tumor samples for the development of the patient-derived xenografts used in this study. Their invaluable contributions have made this research possible.

## Institutional Review Board Statement

The animal study protocol was approved by the Institutional Animal Care and Use Committee at West Virginia University (1603001124, date of approval—8 August 2023).

## Data availability

The following data will be made available upon publication. Raw bulk RNA-sequencing data (.fastq) is available for download at BioProject ID: PRJNA1076666 murine samples. The following data are available on Figshare (doi:10.6084/m9.figshare.25458193): raw spectral files (.mzML) from untargeted metabolomics and quantitative lipidomics; raw whole-animal indirect calorimetry files; processed, identified metabolites/lipids. Any other data are available upon request.

## Author Contributions

*Conceptualization:* S.A.C., E.E.P. *Data collection:* S.A.C., A.D.M., M.W., L.E.R., E.M., W.G., E.E.P. *Data analysis and interpretation:* S.A.C., A.D.M., M.W., L.E.R., E.E.P. *Drafting of article:* S.A.C., E.E.P. *Critical revision of article:* S.A.C., A.D.M., M.W., L.E.R., W.G., E.E.P. *Visualization:* S.A.C., A.D.M., L.E.R. *Supervision:* E.E.P. *Project administration:* M.A.W. and E.E.P. *Funding acquisition:* E.E.P. All authors reviewed the results and approved the final version for publication.

