## Supplemental Figures for "Preclinical Multi-Omic Assessment of Pioglitazone in Skeletal Muscles of Mice Implanted with Human HER2/neu Overexpressing Breast Cancer Xenografts"

**Supplementary Figure 1. STR profile comparison of original tumor with passaged injected HER2/neu overexpressing PDX tumor.** (**a**) STR profile of original patient tumor sample. (**b**) STR profile of passaged injected tumor sample.

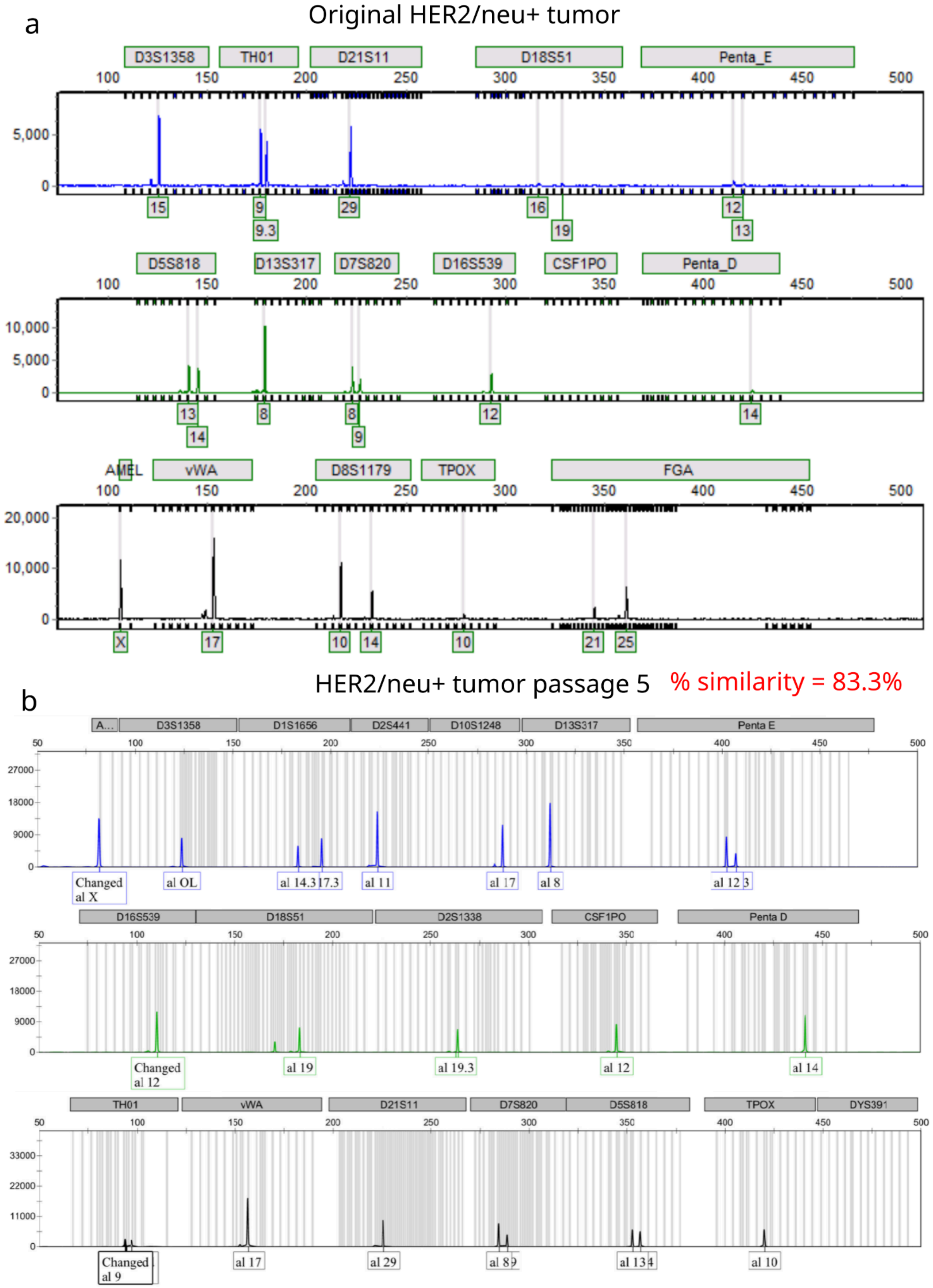

**Supplementary Figure 2. Additional bulk RNA-seq analysis.** (**a**) Bioanalyzer image of all samples (B1-H1, A2-D2). A1 is the ladder. Note: figure includes samples that are not part of this study (E2-H2, A3-C3). (**b**) Total raw read counts in millions for pioglitazone (salmon) and vehicle (blue).. (**c**) Standard deviation of all genes and top 1000 (red dotted line). (**d**) SCREE plot of principal component expected variation. (**e**) Lollipop plot of k-means cluster 5 showing enrichment of PPAR signaling pathway. (**f**) GSEA enrichment of the PPAR signaling pathway. NES = 1.37 and FDR q-value = 0.071.

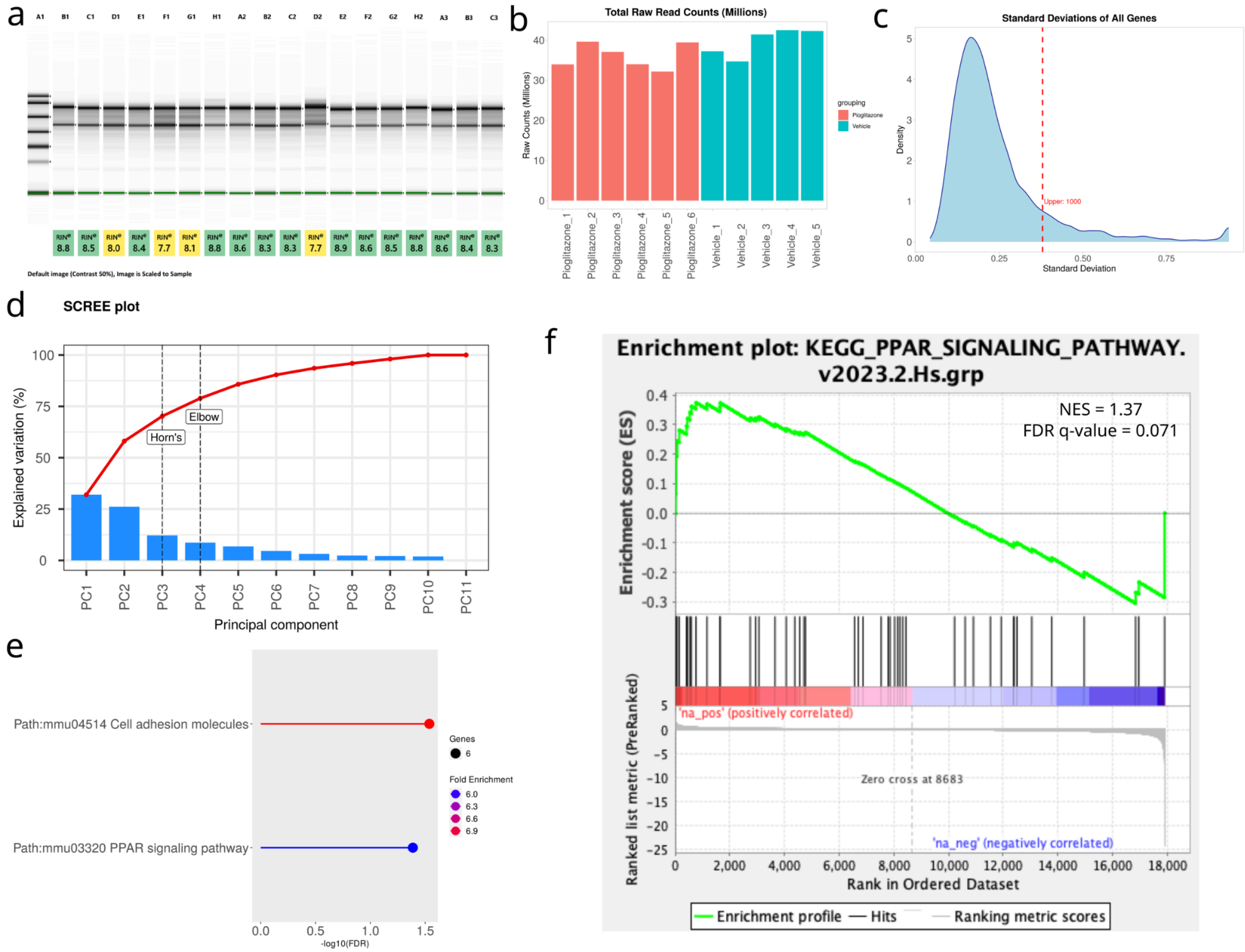

**Supplementary Figure 3. Quality control (QC) metrics for quantitative lipidomics.** (**a**) Ring plot describing proportion of each compound class combined from all samples. (**b**) Pearson correlation analysis of QC samples. The |r| values in upper right squares represent correlation between QC samples, the closer to 1 the better. (**c**) Two-dimensional principal component analysis (PCA) plot including QC samples. (**d**) Coefficient of variation (CV) plot for QC and sample groups. Horizontal axis represents CV value and vertical axis represents the percent of peaks (proportion of metabolites). (**e**) Principal component 1 (PC1) variation of all samples. Horizontal axis represents injection order of samples and vertical axis represents standard deviation (SD) of PC1 score.

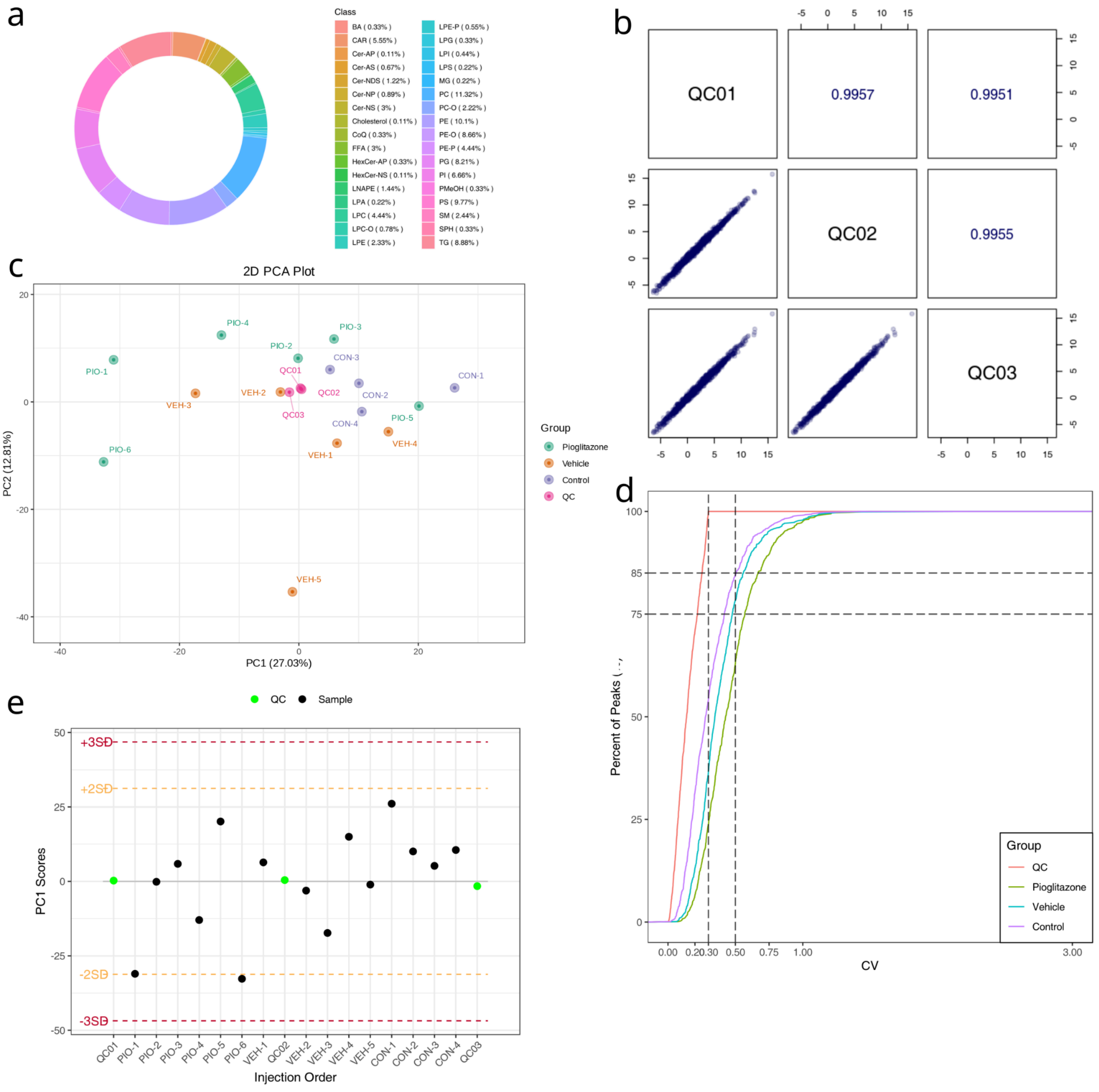

**Supplementary Figure 4. Lipidomics OPLS-DA model validations and PCA plots.** (**a-c**) Orthogonal partial least squares discriminant analysis (OPLS-DA) model validation for pioglitazone vs. vehicle (**a**), vehicle vs. control (**b**), and pioglitazone vs. control (**c**). (**d-f**) 2D PCA plot of pioglitazone (green) vs. vehicle (orange) (**d**), vehicle (green) vs. control (orange) (**e**), and pioglitazone (green) vs. control (orange) (**f**).

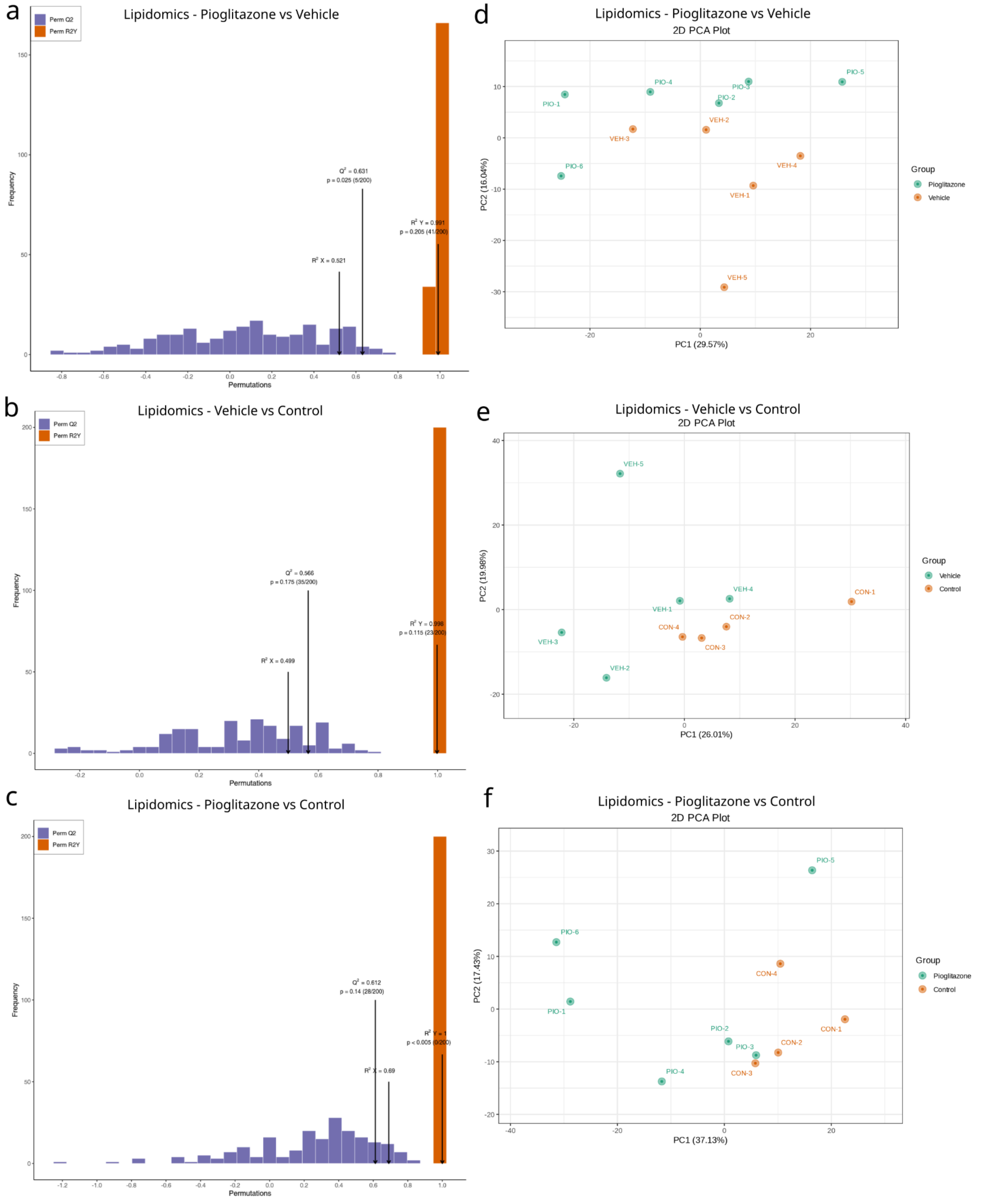

**Supplementary Figure 5. Quality control (QC) metrics for untargeted metabolomics.** (**a**) Ring plot describing proportion of each compound class combined from all samples. (**b**) Pearson correlation analysis of QC samples. The |r| values in upper right squares represent correlation between QC samples, the closer to 1 the better. (**c**) Two-dimensional principal component analysis (PCA) plot including QC samples. (**d**) Coefficient of variation (CV) plot for QC and sample groups. Horizontal axis represents CV value and vertical axis represents the percent of peaks (proportion of metabolites). (**e**) Principal component 1 (PC1) variation of all samples. Horizontal axis represents injection order of samples and vertical axis represents standard deviation (SD) of PC1 score.

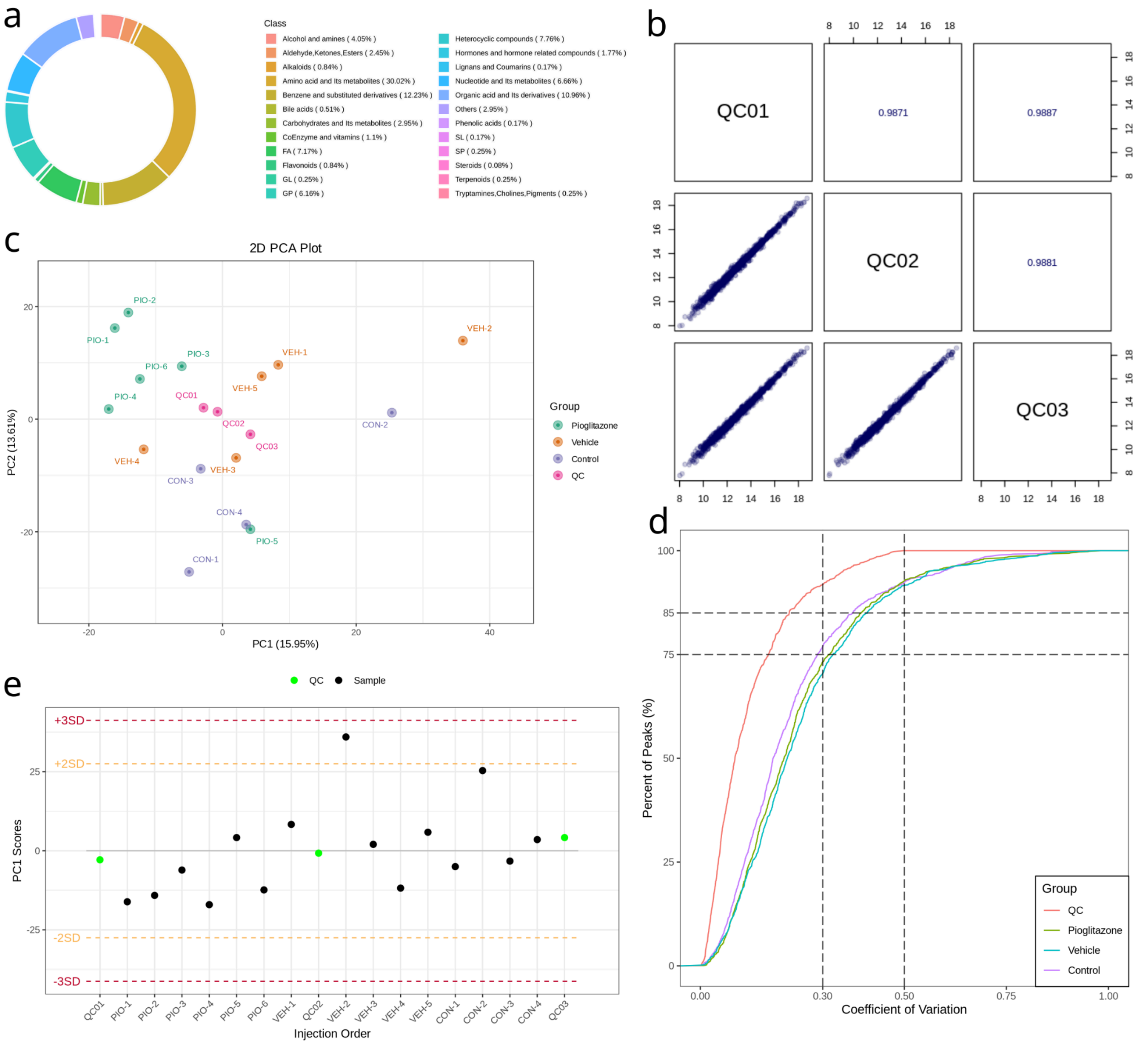

**Supplementary Figure 6. Metabolomics OPLS-DA model validations and PCA plots.** (**a-c**) Orthogonal partial least squares discriminant analysis (OPLS-DA) model validation for pioglitazone vs. vehicle (**a**), vehicle vs. control (**b**), and pioglitazone vs. control (**c**). (**d-f**) 2D PCA plot of pioglitazone (green) vs. vehicle (orange) (**d**), vehicle (green) vs. control (orange) (**e**), and pioglitazone (green) vs. control (orange) (**f**).

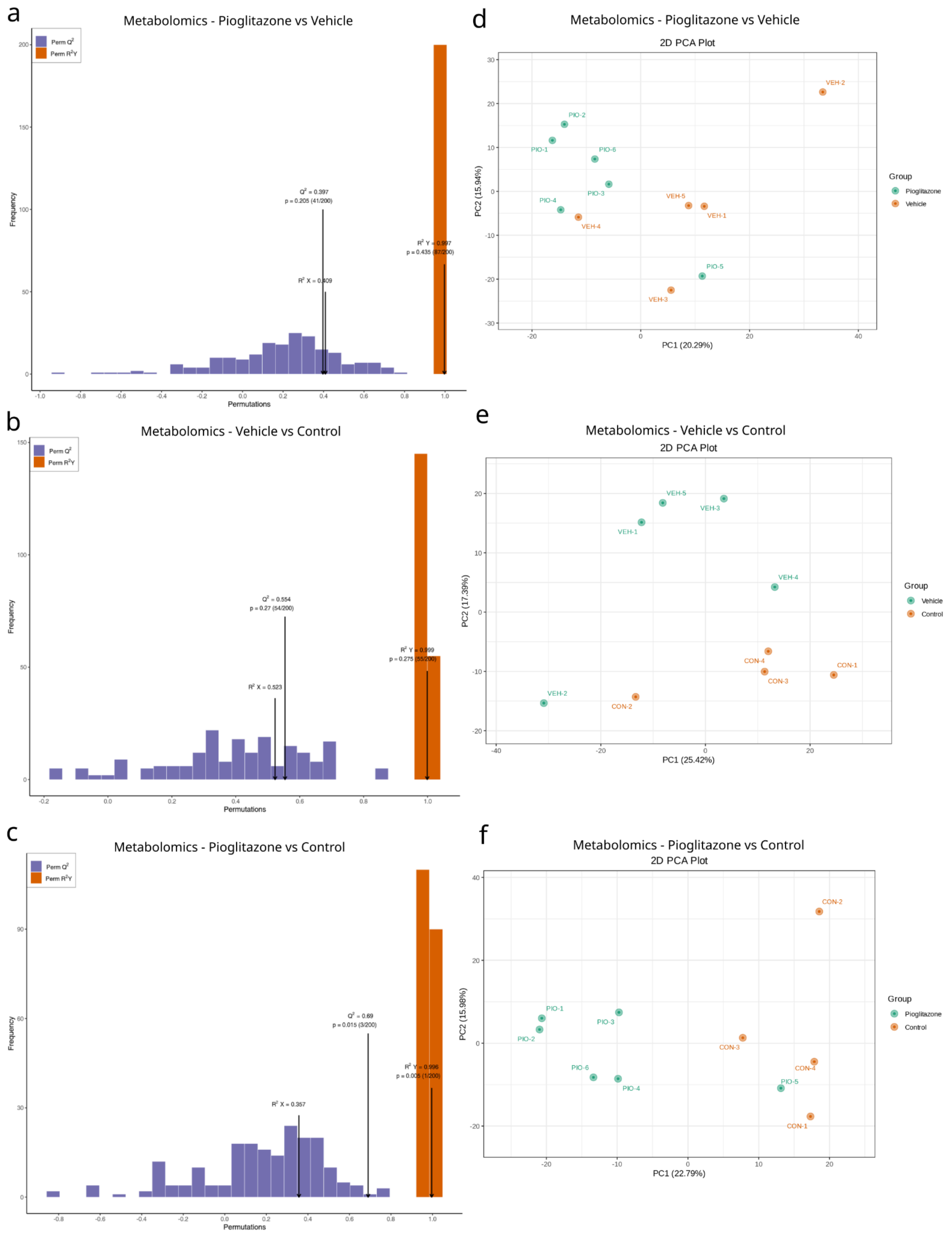

**Supplementary Figure 7. Additional pathways from untargeted metabolomics for vehicle vs. control (left side) and PioTx vs. vehicle (right side).** Asterisks indicate FDR-corrected p values < 0.05. (**a**) Metabolite set enrichment analysis (MSEA) of vehicle vs. control. (**b**) MSEA of PioTx vs. vehicle. No FDR-corrected significant *p*-values.

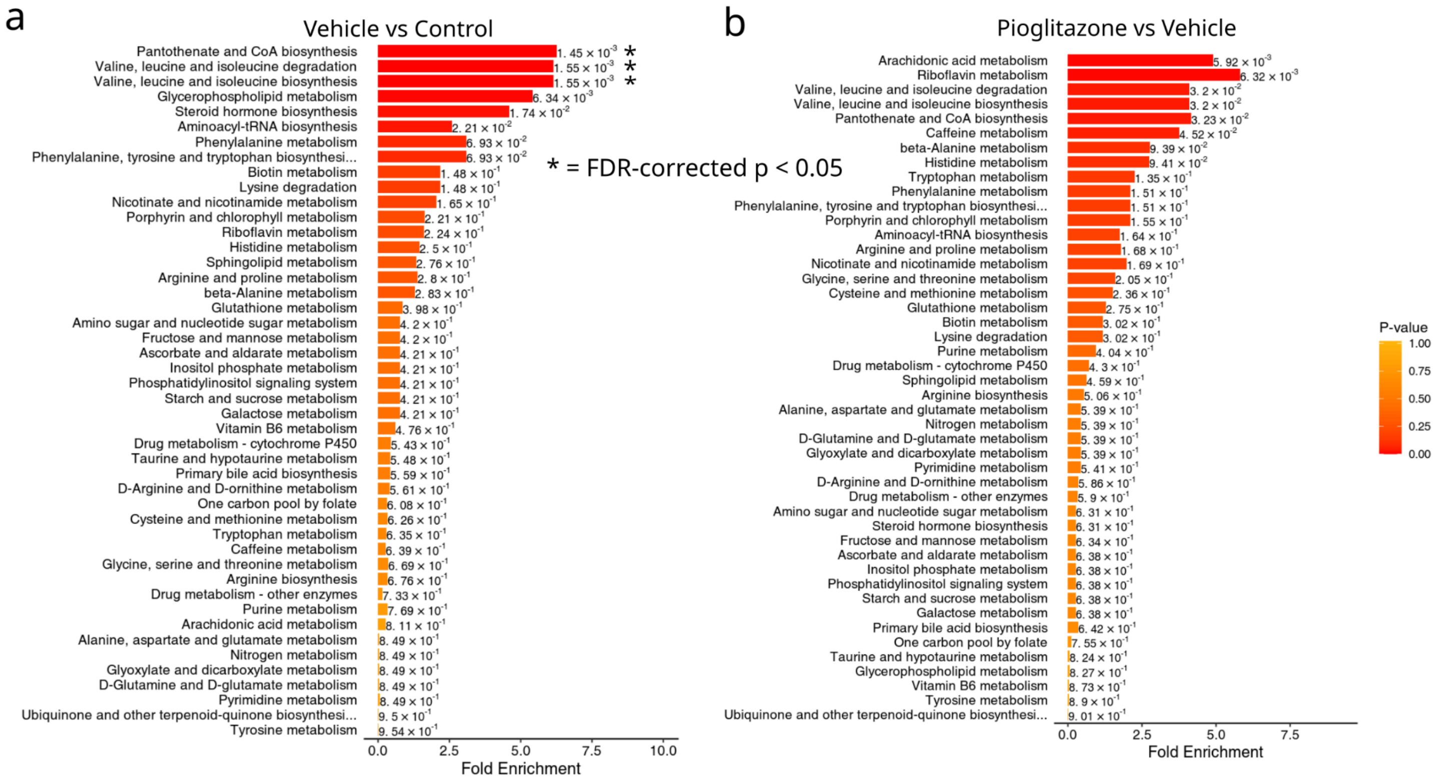

**Supplementary Figure 8. *Ex vivo* soleus functional testing.** (**a**) Two-way ANOVA analysis of all soleus contractions normalized to first repetition. Plotted as first repetition and every subsequent tenth repetition using normalized data points. (**b**) Area under the fatigue curve (AUC) for data presented in **a**. (**c**) Extensor digitorum longus (EDL) absolute force-frequency relationship (FFR) curve. (**d**) EDL normalized FFR curve. (**e**) Soleus absolute FFR curve. (**f**) Soleus normalized FFR curve. (**a-f**) Black dotted lines represent mean values for a naive NSG mouse. Error bars represent the standard error of the mean (SE). (**d, f**) Muscle force normalized to percent of maximal theoretical force as determined by the $P_{max}$ parameter for each individual animal's FFR regression model.

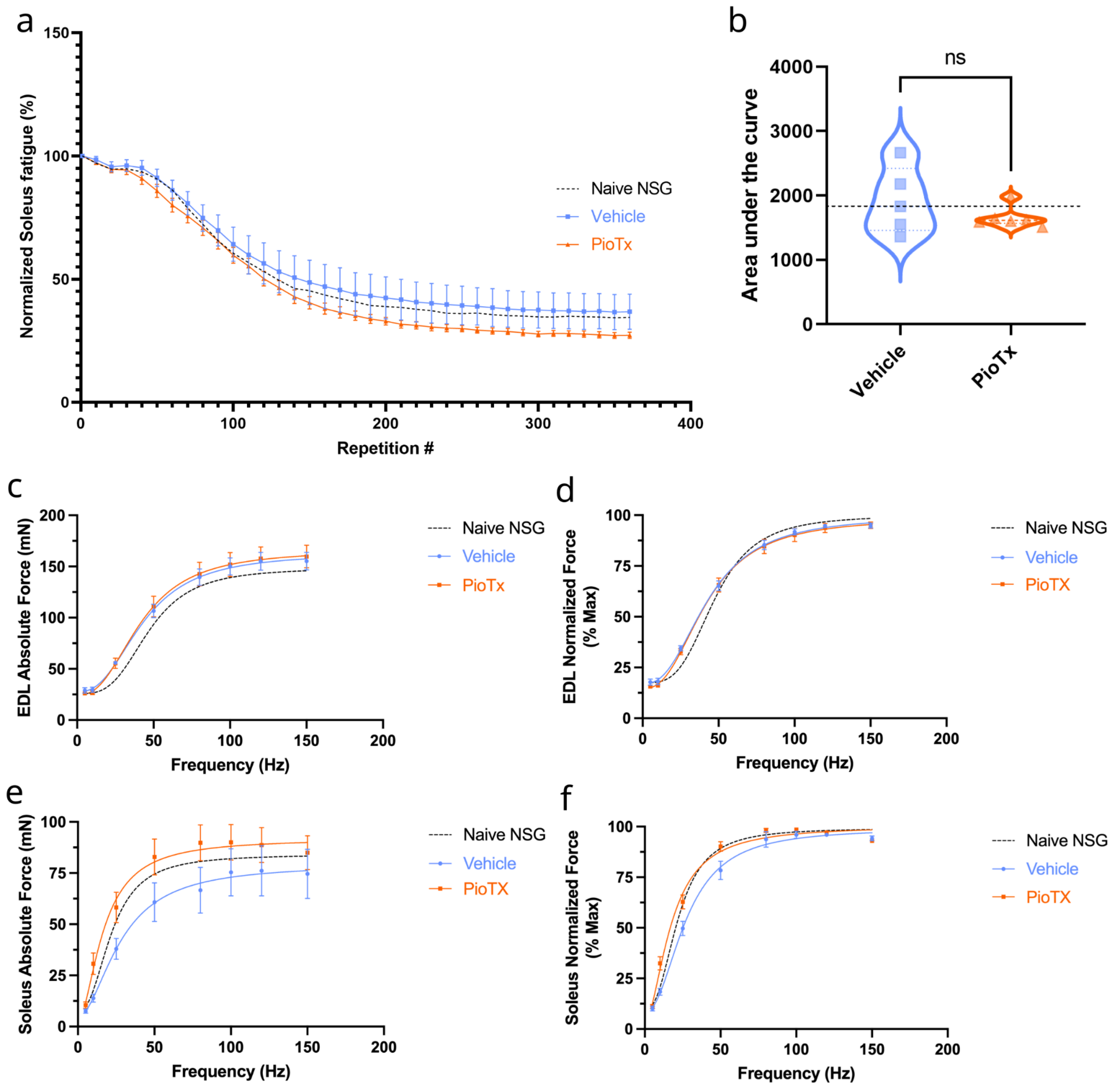
