## Supplemental Tables for "Preclinical Multi-Omic Assessment of Pioglitazone in Skeletal Muscles of Mice Implanted with Human HER2/neu Overexpressing Breast Cancer Xenografts"

**Supplementary Table 1.** Sample sizes used for all components of study. For each group, values indicate the number of samples included in analysis. PioTx = pioglitazone-treated.

| | Variable | Study Group | | |
|---|---|---|---|---|
| | | PioTx | Vehicle | Naive NSG |
| Metabolomics/Lipidomics | All | 6 | 5 | 4 |
| Whole-animal indirect calorimetry | All | 5 | 4 | |
| Bulk RNA-seq | All | 6 | 5 | |
| Weights | TA | 12 | 9 | |
| | EDL | 12 | 9 | |
| | Gastrocnemius | 12 | 9 | |
| | Soleus | 12 | 9 | |
| | Body Mass | 6 | 5 | |
| Muscle Contractile Properties | $L_o$ | 6 | 5 | |
| | CSA | 6 | 5 | |
| | Peak Twitch | 6 | 5 | |
| | Peak CT | 6 | 5 | |
| | Peak RFD | 6 | 5 | |
| | Peak ½ RT | 6 | 5 | |
| | Peak RR | 6 | 5 | |
| | Peak Tetanus | 6 | 5 | |
| | Twitch/CSA | 6 | 5 | |
| | Tetanus/CSA | 6 | 5 | |
| | Twitch/Tetanus | 6 | 5 | |
| Force-Frequency | Force (@ each frequency) | 6 | 5 | |
| Fatigue | Peak Force per Rep | 6 | 5 | |

**Supplementary Table 2.** *Ex vivo* isometric contractile metrics for EDL. Values reported as mean ± SD. $L_o$ = optimal length, CSA = cross-sectional area, mN = milliNewton.

| EDL Isometrics | | | |
|---|---|---|---|
| | **Vehicle** | **Pioglitazone** | ***p*-value** |
| EDL $L_o$ (mm) | 10.3 ± 0.7 | 10.6 ± 0.5 | 0.4951 |
| EDL CSA ($mm^2$) | 1.2 ± 0.3 | 1.4 ± 0.2 | 0.2583 |
| Twitch (mN) | 28.6 ± 7.2 | 25.5 ± 4.3 | 0.4013 |
| Twitch (mN * $CSA^{-1}$) | 25.0 ± 6.4 | 18.8 ± 4.3 | 0.0871 |
| Contraction time (ms) | 20.0 ± 7.1 | 23.3 ± 8.2 | 0.4927 |
| Rate of force development (mN * $s^{-1}$) | 1082 ± 318.3 | 910.5 ± 116.6 | 0.2478 |
| ½ Relaxation time (ms) | 28.0 ± 4.5 | 31.7 ± 4.1 | 0.1889 |
| Rate of relaxation (mN * $s^{-1}$) | -702.7 ± 216.9 | -555.5 ± 144.1 | 0.2101 |
| Tetanus (mN) | 156.2 ± 19.4 | 160.0 ± 27.0 | 0.7968 |
| Tetanus (mN * $CSA^{-1}$) | 136.8 ± 24.4 | 117.3 ± 24.5 | 0.2192 |
| $P_{min}$ (mN) | 27.62 ± 5.249 | 24.48 ± 7.375 | 0.7387 |
| $P_{max}$ (mN) | 163.0 ± 8.140 | 165.8 ±10.89 | 0.8393 |
| $K_f$ (Hz) | 43.12 ± 3.852 | 41.71 ± 4.941 | 0.8308 |
| Hill Slope | 2.535 ± 0.5380 | 2.525 ± 0.7068 | 0.9914 |

**Supplementary Table 3.** *Ex vivo* isometric contractile metrics for soleus. Values reported as mean ± SD. $L_o$ = optimal length, CSA = cross-sectional area, mN = milliNewton.

| Soleus Isometrics | | | |
|---|---|---|---|
| | **Vehicle** | **Pioglitazone** | ***p*-value** |
| Soleus $L_o$ (mm) | 8.5 ± 0.5 | 8.8 ± 0.7 | 0.4311 |
| Soleus CSA ($mm^2$) | 1.0 ± 0.3 | 0.9 ± 0.3 | 0.7512 |
| Twitch (mN) | 7.3 ± 2.3 | 9.9 ± 3.7 | 0.2096 |
| Twitch (mN * $CSA^{-1}$) | 8.9 ± 6.3 | 12.4 ± 5.9 | 0.3700 |
| Contraction time (ms) | 52.0 ± 11.0 | 61.7 ± 11.7 | 0.1938 |
| Rate of force development (mN * $s^{-1}$) | 135.8 ± 49.3 | 151.8 ± 37.0 | 0.5546 |
| ½ Relaxation time (ms) | 74.0 ± 28.8 | 106.7 ± 19.7 | 0.0524 |
| Rate of relaxation (mN * $s^{-1}$) | -51.9 ± 53.3 | -135.4 ± 210.4 | 0.4135 |
| Tetanus (mN) | 76.6 ± 26.7 | 90.4 ± 21.4 | 0.3652 |
| Tetanus (mN * $CSA^{-1}$) | 90.0 ± 55.0 | 109.3 ± 39.3 | 0.5127 |
| $P_{min}$ (mN) | 4.654 ± 13.26 | 2.233 ± 21.03 | 0.9228 |
| $P_{max}$ (mN) | 79.28 ± 12.04 | 91.78 ± 7.274 | 0.4481 |
| $K_f$ (Hz) | 28.12 ± 8.686 | 16.76 ± 5.934 | 0.2567 |
| Hill Slope | 1.876 ± 1.295 | 1.763 ± 0.9354 | 0.9404 |

**Supplementary Table 4.** Top 5 up- and downregulated pathways in PioTx vs. vehicle for all gene set sources. All adjusted (Adj.) *p*-value significant pathways can be found in Supplementary Data 5-9.

| **PioTx vs. Vehicle** | | | | | |
|---|---|---|---|---|---|
| Source | Direction | Pathway | Statistic | # of genes | Adj. Pval |
| GO Bio | Down | GO:0030334 regulation of cell migration | -4.6764 | 491 | 0.006 |
| | | GO:0061061 muscle structure development | -4.5051 | 420 | 0.0069 |
| | | GO:0060537 muscle tissue development | -4.3546 | 257 | 0.01 |
| | | GO:0030335 positive regulation of cell migration | -4.177 | 294 | 0.014 |
| | | GO:0040017 positive regulation of locomotion | -4.121 | 306 | 0.014 |
| | Up | GO:0042776 proton motive force-driven mitochondrial ATP synthesis | 7.2518 | 61 | < 0.001 |
| | | GO:0042773 ATP synthesis coupled electron transport | 7.2144 | 70 | < 0.001 |
| | | GO:0022904 respiratory electron transport chain | 7.0116 | 86 | < 0.001 |
| | | GO:0042775 mitochondrial ATP synthesis coupled electron transport | 7.0104 | 68 | < 0.001 |
| | | GO:0009060 aerobic respiration | 6.9121 | 162 | < 0.001 |
| GO Cell | Down | GO:0031252 cell leading edge | -3.8918 | 252 | 0.016 |
| | | GO:0015629 actin cytoskeleton | -3.6586 | 289 | 0.016 |
| | | GO:0099512 supramolecular fiber | -3.6304 | 488 | 0.016 |
| | | GO:0099081 supramolecular polymer | -3.6295 | 490 | 0.016 |
| | | GO:0042641 actomyosin | -3.393 | 61 | 0.041 |
| | Up | GO:0098798 mitochondrial protein-containing complex | 10.7536 | 272 | < 0.001 |
| | | GO:0005743 mitochondrial inner membrane | 9.7133 | 418 | < 0.001 |
| | | GO:0098800 inner mitochondrial membrane protein complex | 9.4954 | 137 | < 0.001 |
| | | GO:0019866 organelle inner membrane | 9.0888 | 446 | < 0.001 |
| | | GO:0070469 respirasome | 8.3884 | 88 | < 0.001 |
| GO Mol | Down | GO:0003779 actin binding | -3.9998 | 257 | 0.019 |
| | | GO:0008134 transcription factor binding | -3.3466 | 405 | 0.049 |
| | | GO:0019900 kinase binding | -3.3391 | 491 | 0.049 |
| | | GO:0061629 RNA polymerase II-specific DNA-binding transcription factor binding | -3.3022 | 249 | 0.049 |
| | | GO:0019901 protein kinase binding | -3.201 | 439 | 0.049 |
| | Up | GO:0016491 oxidoreductase activity | 5.1059 | 358 | < 0.001 |
| | | GO:0015453 oxidoreduction-driven active transmembrane transporter activity | 4.867 | 28 | 0.002 |
| | | GO:0009055 electron transfer activity | 4.5014 | 49 | 0.002 |
| Reactome | Up | R-MMU-163200 Respiratory electron transport ATP synthesis by chemiosmotic coupling and heat production by uncoupling proteins | 8.2408 | 113 | < 0.001 |
| | | R-MMU-1428517 The citric acid TCA cycle and respiratory electron transport | 7.9483 | 154 | < 0.001 |
| | | R-MMU-611105 Respiratory electron transport | 7.4995 | 94 | < 0.001 |
| | | R-MMU-6799198 Complex I biogenesis | 6.5292 | 54 | < 0.001 |
| | | R-MMU-5368287 Mitochondrial translation | 5.3255 | 88 | < 0.001 |
| KEGG | Up | Path:mmu00190 Oxidative phosphorylation | 7.4728 | 114 | < 0.001 |
| | | Path:mmu04714 Thermogenesis | 5.294 | 187 | < 0.001 |
| | | Path:mmu05415 Diabetic cardiomyopathy | 4.7908 | 160 | < 0.001 |
| | | Path:mmu05208 Chemical carcinogenesis-reactive oxygen species | 4.0763 | 167 | 0.002 |
| | | Path:mmu04723 Retrograde endocannabinoid signaling | 4.0202 | 76 | 0.0026 |

**Supplementary Table 5.** Abbreviations and corresponding complete names for all identified lipid classes.

| | | | | | |
|---|---|---|---|---|---|
| **BA** | bile acids | **CAR** | carnitines | **Cer-AP** | ceramide alpha-hydroxy fatty acid-phytosphingosine |
| **Cer-AS** | ceramide alpha-hydroxy fatty acid-sphingosine | **Cer-NDS** | ceramide non-hydroxy fatty acid-dihydrosphingosine | **Cer-NP** | ceramide non-hydroxy fatty acid-phytosphingosine |
| **Cer-NS** | ceramide alpha-hydroxy fatty acid-sphingosine | **Cholesterol** | cholesterol | **CoQ** | coenzyme Q |
| **FFA** | free fatty-acids | **HexCer-AP** | hexosylceramide alpha-hydroxy fatty acid-phytosphingosine | **HexCer-NS** | Hexosylceramide non-hydroxy fatty acid-sphingosine |
| **LNAPE** | N-acyl-lysophosphatidylethan olamine | **LPA** | lysophosphatidic acid | **LPC** | lysophosphatidylcholine |
| **LPC-O** | alkyl-lysophosphatidylcholine | **LPE** | lysophosphatidylethanolamine | **LPE-P** | alkenyl-lysophosphatidylethan olamine |
| **LPG** | lysophosphatidylglycerol | **LPI** | lysophosphatidylinositol | **LPS** | lysophosphatidylserine |
| **MG** | monoacylglycerol | **PC** | phosphatidylcholine | **PC-O** | alkyl-phosphatidylcholine |
| **PE** | phosphatidylethanolamine | **PE-O** | alkyl-phosphatidylethanolamine | **PE-P** | alkenyl-phosphatidylethanola mine |
| **PG** | phosphatidylglycerol | **PI** | phosphatidylinositol | **PMeOH** | phosphatidylmethanol |
| **PS** | phosphatidylserine | **SM** | sphingomyelin | **SPH** | sphingosine |
| **TG** | triacylglycerol | | | | |

**Supplementary Table 6.** Weight in milligrams of all isolated muscles from all groups of mice. Muscles isolated were extensor digitorum longus (EDL), soleus, tibialis anterior (TA), and gastrocnemius (Gastroc). Missing TA length values and corresponding normalizations were due to fractured tibias and not included in analysis. TL: tibia length (mm). / TL: muscle weight divided by tibia length. PIO: tumor-bearing and pioglitazone treated. VEH: tumor-bearing and vehicle treated.

| | **ID** | **Limb** | **TL (mm)** | **EDL** | | **Soleus** | | **TA** | | **Gastroc** | |
|---|---|---|---|---|---|---|---|---|---|---|---|
| | | | | **mg** | **/ TL** | **mg** | **/ TL** | **mg** | **/ TL** | **mg** | **/ TL** |
| PIO | PIO-1 | R | 17.84 | 8 | 0.448 | 7 | 0.392 | 43 | 2.410 | 118 | 6.614 |
| | | L | 17.77 | 10 | 0.563 | 8 | 0.450 | 44 | 2.476 | 139 | 7.822 |
| | PIO-2 | R | 18.1 | 7 | 0.387 | 6 | 0.331 | 43 | 2.376 | 126 | 6.961 |
| | | L | 18.4 | 9 | 0.489 | 7 | 0.380 | 43 | 2.337 | 156 | 8.478 |
| | PIO-3 | R | 18.26 | 7 | 0.383 | 6 | 0.329 | 49 | 2.683 | 143 | 7.831 |
| | | L | 18.33 | 9 | 0.491 | 8 | 0.436 | 47 | 2.564 | 156 | 8.511 |
| | PIO-4 | R | 18.45 | 7 | 0.379 | 8 | 0.434 | 47 | 2.547 | 118 | 6.396 |
| | | L | 18.43 | 8 | 0.434 | 10 | 0.543 | 45 | 2.442 | 88 | 4.775 |
| | PIO-5 | R | 18.27 | 8 | 0.438 | 7 | 0.383 | 49 | 2.682 | 122 | 6.678 |
| | | L | 18.32 | 10 | 0.546 | 3 | 0.164 | 47 | 2.566 | 173 | 9.443 |
| | PIO-6 | R | 17.8 | 5 | 0.281 | 4 | 0.225 | 32 | 1.798 | 126 | 7.079 |
| | | L | 18.9 | 7 | 0.370 | 8 | 0.423 | 35 | 1.852 | 109 | 5.767 |
| VEH | VEH-1 | R | 17.54 | 6 | 0.342 | 9 | 0.513 | 39 | 2.223 | 114 | 6.499 |
| | | L | 17.06 | 6 | 0.352 | 11 | 0.645 | 38 | 2.227 | 119 | 6.975 |
| | VEH-2 | R | 17.2 | 5 | 0.291 | 7 | 0.407 | 46 | 2.674 | 144 | 8.372 |
| | | L | / | 5 | / | 13 | / | 49 | / | 146 | / |
| | VEH-3 | R | 18.34 | 8 | 0.436 | 5 | 0.273 | 41 | 2.236 | 127 | 6.925 |
| | | L | 18.51 | 8 | 0.432 | 9 | 0.486 | 39 | 2.107 | 130 | 7.023 |
| | VEH-4 | R | 18.14 | 6 | 0.331 | 6 | 0.331 | 41 | 2.260 | 140 | 7.718 |
| | | L | 18.18 | 11 | 0.605 | 10 | 0.550 | 41 | 2.255 | 141 | 7.756 |
| | VEH-5 | R | 18.0 | 4 | 0.222 | 4 | 0.222 | 33 | 1.833 | 107 | 5.944 |
| | | L | 17.87 | 6 | 0.336 | 6 | 0.336 | 29 | 1.623 | 106 | 5.932 |
