## Supplemental Data List for "Preclinical Multi-Omic Assessment of Pioglitazone in Skeletal Muscles of Mice Implanted with Human HER2/neu Overexpressing Breast Cancer Xenografts"

**Supplementary Data 1.** Untargeted metabolomics internal standard stability in QC samples.

**Supplementary Data 2.** Cumulative mouse body weights in grams.

**Supplementary Data 3.** Unnormalized gene expression counts for Pioglitazone and Vehicle groups.

**Supplementary Data 4.** K-means cluster-5 gene list.

**Supplementary Data 5.** Pioglitazone- vs. vehicle-treated all adjusted *p*-value significant pathways from KEGG.

**Supplementary Data 6.** Pioglitazone- vs. vehicle-treated all adjusted *p*-value significant pathways from GO Biological.

**Supplementary Data 7.** Pioglitazone- vs. vehicle-treated all adjusted *p*-value significant pathways from GO Cellular.

**Supplementary Data 8.** Pioglitazone- vs. vehicle-treated all adjusted *p*-value significant pathways from GO Molecular.

**Supplementary Data 9.** Pioglitazone- vs. vehicle-treated all adjusted *p*-value significant pathways from Reactome.
